# Nanopore adaptive sampling to identify the NLR-gene family in melon (*Cucumis melo* L.)

**DOI:** 10.1101/2023.12.20.572599

**Authors:** Javier Belinchon-Moreno, Aurelie Berard, Aurelie Canaguier, Véronique Chovelon, Corinne Cruaud, Stéfan Engelen, Rafael Feriche-Linares, Isabelle Le-Clainche, William Marande, Vincent Rittener-Ruff, Jacques Lagnel, Damien Hinsinger, Nathalie Boissot, Patricia Faivre Rampant

## Abstract

**Background:** Nanopore Adaptive Sampling (NAS) offers a promising approach for assessing genetic diversity in targeted genomic regions. Herein, we design and validate an experiment to enrich a set of resistance genes in several melon cultivars as a proof of concept.

**Results:** We showed that each of the 15 regions we identified in two newly assembled melon genomes (subspecies *melo*) were successfully and accurately reconstructed as well as in a third cultivar from the *agrestis* subspecies. We obtained a fourfold enrichment, independently from the samples, but with some variations according to the enriched regions. In the *agrestis* cultivar, we further confirmed our assembly by PCR. We discussed parameters that can influence enrichment and accuracy of assemblies generated through NAS.

**Conclusions:** Altogether, we demonstrated NAS as a simple and efficient approach to explore complex genomic regions. This approach finally unlocks the characterization of resistance genes for a large number of individuals, as required for breeding new cultivars responding to the agroecological transition.

## Background

The correct assembly of complex and highly repeated genome regions, especially notorious in plants, remains a challenge. Despite being widely used on whole genome sequencing (WGS) approaches due to their cost-effectiveness, short-read sequencing methods prove ineffective in these complex regions as their length cannot allow a proper assessment of copy-number variation and duplication events (Lee & Chae, 2020). Long-read technologies, such as those provided by Oxford Nanopore Technologies (ONT, Oxford, UK), demonstrated their potential in accurately resolving complex regions by spanning long repetitive elements or areas with tandemly repeated genes (Mohamed et al., 2020; Lieberman et al., 2022). Nevertheless, whole-genome long-read sequencing remains cost-ineffective for most studies, particularly those requiring the sequencing of numerous genotypes for a few specific regions of interest.

Targeted sequencing approaches are a valuable alternative for characterizing specific genomic regions while reducing sequencing and data storage costs compared to WGS (Hook & Timp, 2023). Current targeted sequencing protocols have been adapted for long-read sequencing and predominantly employ hybridization capture (Witek et al., 2016), PCR amplification ((Norris et al., 2016), Cas9-assisted targeting methods (Gilpatrick et al., 2020), or microfluidic-based droplet sorting procedures (Madsen et al., 2020). However, these approaches require substantial experimental and design efforts, along with a high prior knowledge of the sequence to be enriched and its genotypic diversity (Gilpatrick et al., 2020; Hook & Timp, 2023). Particularly, PCR-based techniques are prone to introducing biases in the enriched sequences, and long amplicons are difficult to consistently amplify (Hook & Timp, 2023). Hybridization-based methods require the construction of complex RNA libraries together with very specific hybridization and capture conditions (Hook & Timp, 2023). Cas9-based methods like Nanopore Cas9-targeted sequencing (nCATS) (Gilpatrick et al., 2020) or Cas9-Assisted Targeting of Chromosome segments (CATCH) (Gabrieli et al., 2017) require the design of multiple guide-RNAs, a task that may be challenging for complex and repetitive genome regions. Finally, microfluidic-based methods like Xdrop (Madsen et al., 2020) are highly complex and require specialized microfluidic equipment (Hook & Timp, 2023).

The Nanopore Adaptive Sampling (NAS) approach recently implemented by ONT overcome these limitations. NAS was first suggested in 2016 (Loose et al., 2016) and has been implemented with different algorithms since the end of 2019 (Edwards et al., 2019; Kovaka et al., 2021; Payne, Holmes, et al., 2021; Weilguny et al., 2023). It takes advantage of the ability of the pores to control the directional flow of the DNA strand that is being sequenced depending on the applied current’s polarity. By combining live calling of sequenced bases with real-time mapping to a set of DNA sequences provided by the user for enrichment, the DNA strand is dynamically either discarded or fully sequenced based on the similarity of its initial first few hundred bases to the provided reference (Loose et al., 2016). NAS requires a standard library preparation, eliminates the need for DNA amplification, circumvents laborious or expensive experimental design or probes synthesis, and offers real-time selective enrichment (Martin et al., 2022; Miyatake et al., 2022). NAS has been used in clinical settings and for the enrichment of metagenomic samples (Kipp et al., 2021; De Meulenaere et al., 2022; Martin et al., 2022; Greer et al., 2023; Hewel et al., 2023; Su et al., 2023; Wrenn & Drown, 2023). Therefore, NAS emerges as a promising approach for studying target regions, especially those that are highly complex, such as disease-associated repeat loci in humans (Miyatake et al., 2022; Stevanovski et al., 2022).

In plants, immunity is encoded by resistance genes (R genes), frequently organized in complex regions (Liu et al., 2007). Among R genes, Nucleotide-binding site leucine-rich repeat resistance genes (NLRs) form the largest family (Barragan & Weigel, 2021). These genes encode intracellular receptors that play a central role in the so-called effector-triggered immunity (ETI) against pathogens. NLR genes exhibit a highly conserved structure with three main domains (Barragan & Weigel, 2021; Zhang et al., 2022): the N-terminal domain, the central domain, and the C-terminal domain. The N-terminal domain can be a Toll/Interleukin-1 receptor (TIR), a Coiled-coil (CC), or a resistance to Powdery Mildew 8-like (RPW8) domain. The central domain, the most conserved one, is a nucleotide-binding adaptor (NB-ARC), also named as NBS (nucleotide-binding site) domain. This domain plays a crucial role in signal transduction. Finally, the C-terminal domain is often composed of leucine-rich repeats (LRR) with ligand-binding functions. A clustered genomic arrangement is a common characteristic of NLR genes (Van Wersch & Li, 2019). These clusters often result from unequal crossing overs, tandem duplications, or intra-cluster rearrangements (Barragan & Weigel, 2021). In this context, NAS, combining long-read sequencing and target enrichment, should allow the accurate characterization of NLR clusters in plants.

To investigate the ability of NAS to efficiently retrieve the sequence of the complete set of NLR clusters into a species (or NLRome), we selected melon (*Cucumis melo* L.) as a model. Melon genome presents a i/ small genome size; ii/ NLR content estimated at ≈1% of the genome (González et al., 2013) aligning with ONT target size recommendations (Nanopore Community, 2023); and iii/ finely characterized, highly variable, and complex NLR cluster, *Vat* (Chovelon et al., 2021; Boissot et al., 2023), suitable for benchmarking. Among the well-characterized accessions for the *Vat* region, we chose Anso77 (ssp. *melo*) for its highest number of functional *Vat* genes (Chovelon et al., 2021). We also selected Doublon (ssp. *melo*) as an accession with a contrasting *Vat* region structure compared to Anso77 (Chovelon et al., 2021). Therefore, we assembled and annotated their whole genomes and we established Anso77 as the reference to identify the regions of interest (ROIs) for NAS. We assessed the performance of the method in capturing the set of NLR clusters on Anso77 and Doublon. Furthermore, we extended our assessment to an accession belonging to a different subspecies (Chang-Bougi, ssp. *agrestis*) for which a genome was publicly available (Shin et al., 2019).

## Materials and Methods

### BIOLOGICAL MATERIAL

We selected melon cultivars Anso77, Doublon and Chang-Bougi to develop a proof of concept for the NLRome adaptive sampling experiment, with Anso77 serving as the reference. The origin of these cultivars is located in Spain, France and Korea, respectively. Anso77 and Doublon were chosen as sp. *melo* lines belonging to the *inodorus* and *cantalupensis* botanical groups. Chang-Bougi, belonging to the *agrestis* subspecies and specifically to the *makuwa* botanical group, was selected as a distantly related cultivar compared to Anso77 and Doublon. This choice aimed to validate the NAS procedure with cultivars significantly differing from the provided reference. Chang-Bougi belongs to the *agrestis* subspecies and more specifically to the *makuwa* botanical group. Additionally, a draft genome assembly of Chang-Bougi, constructed with Illumina HiSeq reads, was readily available (Shin et al., 2019).

We obtained the seeds from the INRAE Centre for Vegetable Germplasm in Avignon (Salinier et al., 2022) and cultivated them under greenhouse conditions at INRAE GAFL, Avignon, France.

### ANSO77 AND DOUBLON *DE NOVO*WHOLE GENOME SEQUENCING, ASSEMBLY, AND ANNOTATION

We produced whole *de novo* genome assemblies of Anso77 and Doublon by combining long-read sequencing: ONT for Anso77 and ONT combined with Pacific Biosciences (PacBio, Menlo Park, CA, USA) for Doublon. Raw reads were already deposited in the NCBI database under the following Bioproject accession numbers: PRJNA662717 and PRJNA662721 (Chovelon et al., 2021). Bionano optical maps (BioNano Genomics, San Diego, CA, USA); 10x Linked-Reads (Pleasanton, CA, USA) for Anso77; Illumina Novaseq short-read sequencing (Illumina, San Diego, CA, USA); and linkage map information were developed and used to construct genome assemblies. Fully detailed methods and parameters employed for the assemblies and annotations are provided in Additional files: Supplementary Methods.

### NAS ENRICHMENT PANEL DEFINITION AND EXPERIMENTAL DESIGN

We used the cultivar Anso77 as the reference for constructing the target regions for the NAS approach. We predicted the presence of NLR-related genes using NLGenomeSweeper (Toda et al., 2020) with default parameters. This tool approximates the presence of NLR genes by the identification of the well-conserved NBS domain. We defined the regions of interest (ROIs) by grouping predicted NBS domains separated by regions shorter than 1 Mb. To ensure robust read depth coverage on the selected ROIs, we added a 20 kb buffer zone flanking the ROIs to constitute the initial target regions.

We performed a REs annotation within the initial target regions using the CENSOR tool from the curated giri Repbase website (Kohany et al., 2006). Predicted REs longer than 200 bp, as well as sequences shorter than 500 bp located between them, were excluded from the initial target regions. Figure 1 illustrates the definition of the ROIs, target regions, and target regions without REs.

**Figure 1.**
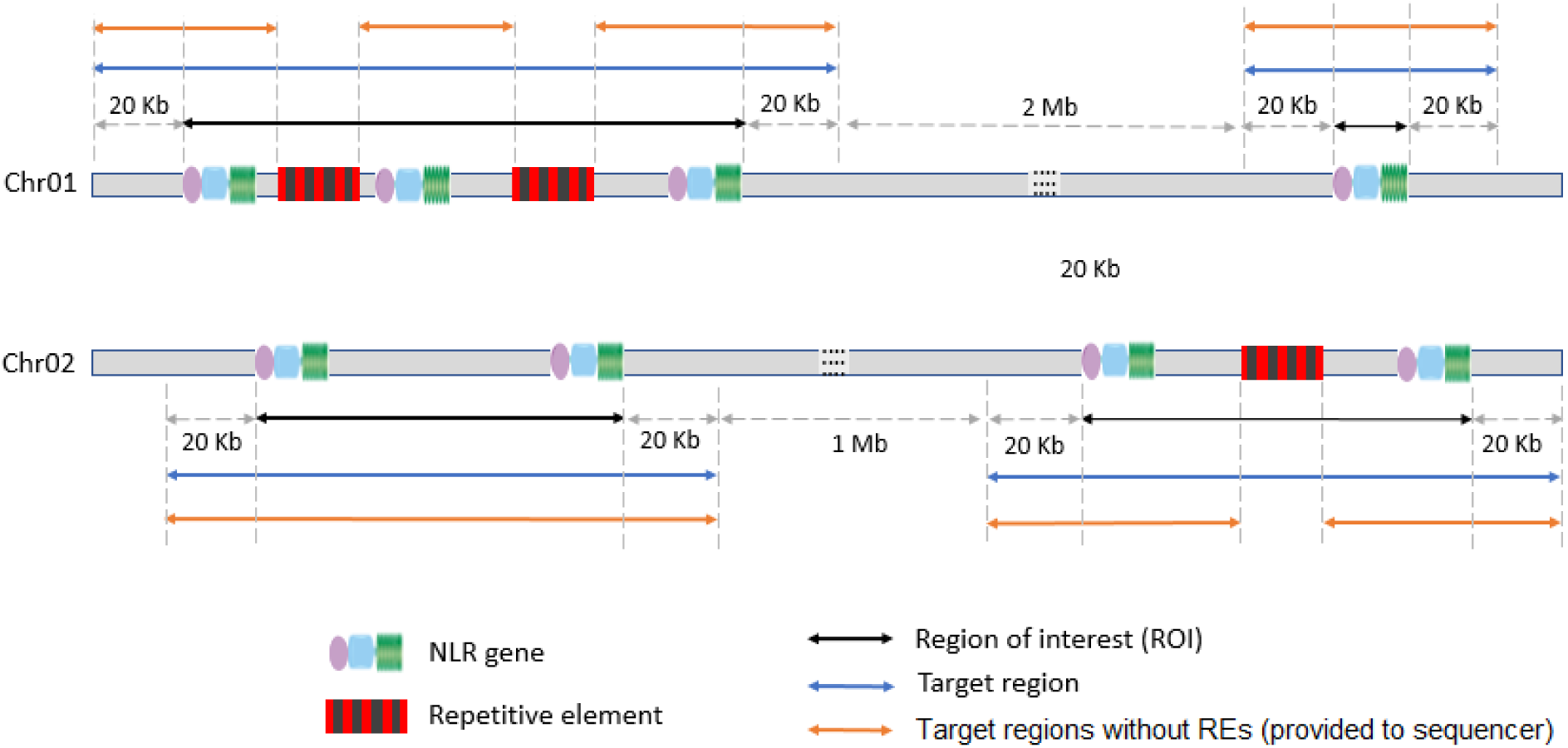
Schematic representation of the definition of ROIs, target regions and target regions without REs (provided to the MinKNOW software) in the reference for NAS, Anso77.

We provided these target regions without REs in bed format and the reference genome of Anso77 in fasta format to the MinKNOW software (ONT, Oxford, UK). These files were used to determine the acceptance or rejection of reads. If the initial ∼500 bps of the DNA strands matched the target regions without REs, they underwent complete sequencing; otherwise, they were rejected from the pore.

### DNA EXTRACTION, ADAPTIVE SAMPLING SEQUENCING, AND BASE-CALLING

Plant leaves were harvested and immediately frozen in liquid nitrogen for subsequent DNA extraction. Genomic DNA was extracted using the NucleoSpin Plant II kit (Macherey-Nagel, Germany) following the manufacturer’s protocol. DNA quantity and quality assessment were conducted using Qubit4® 1x dsDNA BR Assay Kit (Invitrogen, Carlsbad, CA, USA) and Agilent 2200 TapeStation (Agilent Technologies, Santa Clara, CA, USA).

We multiplexed and sequenced DNAs of Anso77 and Doublon on a single PromethION *R10*.4.1 flowcell (ONT, Oxford, UK). Half the channels were used as control channels in which no adaptive sampling was performed. Moreover, we sequenced Chang-Bougi using one-tenth of a PromethION *R10*.4.1 flowcell to assess the flexibility and scalability of NAS. We prepared the sequencing libraries using the Native Barcoding SQK-NBD114.24 (ONT, Oxford, UK) and following ONT guidelines with some modifications. One microgram of genomic DNA from each sample was repaired and end-prepped with an incubation at 20°C for 20 minutes followed by a heat-inactivation of the enzymes at 65 °C for an additional 20 minutes. DNA was purified and barcodes were individually ligated to each of the purified DNA samples. Barcodes NB01 and NB02 were used for Anso77 and Doublon. NB22 was used for Chang-Bougi. Barcoded samples were purified using AMPure XP beads (Beckman Coulter Inc., Brea, CA, USA) at a ratio of 0.4:1 beads-to-barcoding mix, keeping each barcoded sample in an independent Eppendorf tube. Finally, purified barcoded DNA samples were pooled at an equimolar concentration to a total volume of 30 µl. Adapters were ligated to the pooled samples. After purification, DNA size and concentration of the barcoded pools were quantified using Agilent 2200 TapeStation and Qubit4® 1x dsDNA HR Assay Kit (Invitrogen, Carlsbad, CA, USA), respectively. Final libraries were adjusted to a volume of 32 µl containing 10-20 fmol of DNA. All incubations shorter than 10 minutes were extended to 10 minutes.

We completed libraries by adding the Sequencing Buffer (ONT, Oxford, UK) and Library Loading Beads (ONT, Oxford, UK), and subsequently loaded them into R10.4.1 PromethION flowcells for 96-hour runs in the case of Anso77 and Doublon, or 120-hours runs in the case of Chang-Bougi. A library reloading (washing flush) was performed in all the experiences when the percentage of sequencing pores dropped to 10-15%. NAS was performed using channels 1-1500 of the PromethION flowcells for Anso77 and Doublon, keeping the rest of the channels as control. The sequencing speed was set to 260 bps (accuracy mode) for Anso77 and Doublon, and the quality score threshold was set to 10. For Chang-Bougi, NAS was performed on the whole flowcell and the sequenced speed was modified to 400 bps (default mode) because the 260 bps option has been deprecated from MinKNOW version 23.04.

Raw ONT FAST5 files were live base-called during the PromethION run with Guppy (ONT, London, UK) v. 6.3.9 for Anso77 and Doublon and Guppy v. 6.5.7 for Chang-Bougi in “super accurate base-calling” mode. Barcodes were automatically trimmed using the “trim barcodes” option of the MinKNOW software v. 22.10.7 for Anso77 and Doublon, and v. 23.04.5 for Chang-Bougi. For each run, the automatically generated “sequencing_summary.txt” file and the FASTQ files of the samples were retained for further processing.

### NAS DATA PROCESSING AND ENRICHMENT CALCULATION

For Anso77 and Doublon, we split the reads by channel generating two FASTQ files per sample: one with the reads sequenced on channels 1-1500 for NAS, and another with the reads generated on channels 1501-3000 for WGS. No splitting by channel was performed for Chang-Bougi as the totality of the channels were used for NAS. We retained for downstream analyses reads with a “PASS” flag, meaning that their quality score was greater than 10. We identified the rejected reads by NAS based on their “end reason” in the sequencing summary file. This file includes the classification of the generated reads based on their “end reason”. In this way, rejected reads by adaptive sampling are labeled as “Data Service Unblock Mux Change”. Reads labeled as “Unblock Mux Change”, “Mux Change” and “Signal Negative” were also filtered out. Afterwards, we filtered the generated FASTQ files by size, keeping only reads longer than 1 kb. We assessed statistics on these FASTQ files using seqkit stats v. 2.4.0 (Shen et al., 2016).

We computed sequence depth statistics for Anso77 and Doublon by aligning the reads to their reference whole genome assemblies with minimap2 v. 2.24-r1122 (Li, 2018) and using mosdepth v. 0.3.3 (Pedersen & Quinlan, 2018) with a bed file containing the coordinates of the 15 target regions. For Chang-Bougi, sequence depth statistics were calculating by aligning the reads to its assembled target regions. The split flowcell setup allowed the calculation of enrichment in Anso77 and Doublon by comparing read depth generated in NAS and WGS. We assessed the efficiency of NAS using two measures of enrichment. The enrichment by yield, denoted as the ratio of the on-target sequence depth (NLR cluster+20 kb flanking) with NAS to that with WGS, was assessed as follows:

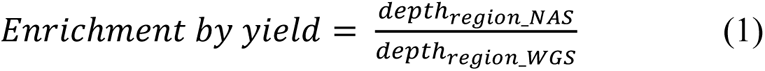

where depth_region_NAS_ and depth_region_WGS_ represent the on-target sequence depth in the NAS and WGS experiments, respectively.

The enrichment by selection, as the ratio of the relative selection of the target regions between NAS and WGS, measures how much NAS can alter the abundance of the given target regions in the context of a complete genome, considering the sequencing behavior of each ROI. It was calculated as follows:

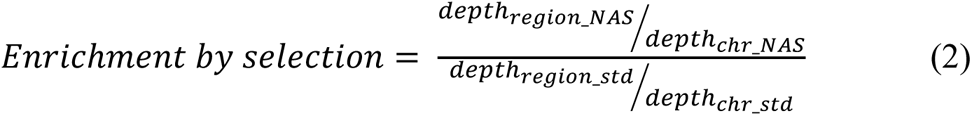

where depth_region_NAS_ and depth_chr_NAS_ represent, respectively, the sequence depth on-target (NLR cluster+20 kb flanking) and on the rest of the chromosome in the adaptive sampling approach, while depth_region_WGS_ and depth_chr_WGS_ represent the depth coverage on-target (NLR cluster+20 kb flanking) and on the rest of the chromosome in the WGS approach. If no bias exists that causes the target regions to be differentially enriched compared to the rest of the genome, the relative selection with WGS should be equal to one.

We calculated average enrichment by yield between all target regions as the ratio of the average sequence depth on-target in NAS and WGS. Similarly, we assessed the average enrichment by selection between all target regions as the ratio of the average relative frequency of target regions in NAS and WGS. The average relative frequencies of target regions were calculated as the ratio of average sequence depth on-target and off-target. Only chromosomes containing target regions were considered in calculating the off-target average sequence depth.

### TARGET REGIONS ASSEMBLY, NLR ANNOTATION AND QUALITY CONTROLS

We tested a set of assemblers tailored for ONT data including Canu (Koren et al., 2017), Flye (Kolmogorov et al., 2019), Shasta (Shafin et al., 2020), Necat (Chen et al., 2021), Raven (Vaser & Šikić, 2021) and SMARTdenovo (Liu et al., 2021), for assembling the NAS reads (data not shown). SMARTdenovo was primarily selected due to its superior assembly metrics (contiguity and assembly errors) of the target regions within a short time and with low memory usage. For Chang-Bougi, one target region was selected from the Canu assembly, as SMARTdenovo failed to collapse a repeat region generating two contigs instead of a single one. We used default parameters and added the “generate consensus” option for SMARTdenovo. Canu was executed with the options “genomesize=7m –corrected –trimmed –nanopore”.

For each assembly, we filtered the contigs retaining only those including at least one predicted NBS domain or matching more than 15 kb with at least 45% identity to any of the 15 target regions of Anso77. We assessed NBS domain prediction using NLGenomeSweeper. We used the nucmer and delta-filter commands from MUMmer’s v. 4.0.0rc1 (Marçais et al., 2018) to select contigs matching the target regions of Anso77. Nucmer was used with the option –l 100, keeping the rest of the parameters as default. Hits reported by nucmer were filtered with delta-filter with the options -r -q −l 15000 -i 45. We ran QUAST v. 5.0.2 to assess the basic statistics of the generated filtered assemblies.

Assembly errors were analysed focusing on the well-studied *Vat* region. We performed manual annotations of the *Vat* regions following (Chovelon et al., 2021). To analyze the accuracy of the *Vat* homologs, we used two PCR markers: Z649 FR which indicates the number of R65aa motifs in the *Vat* homologs (Chovelon et al., 2021), and Z1431 FR which is specific to a *Vat* homolog with four R65aa motifs (Boissot et al., 2023).

Figure 2 depicts a workflow diagram summarizing the different steps involved in data processing and target regions assembly. All statistical tests were performed using R v. 4.1.1 (R Core Team, 2021).

**Figure 2.**
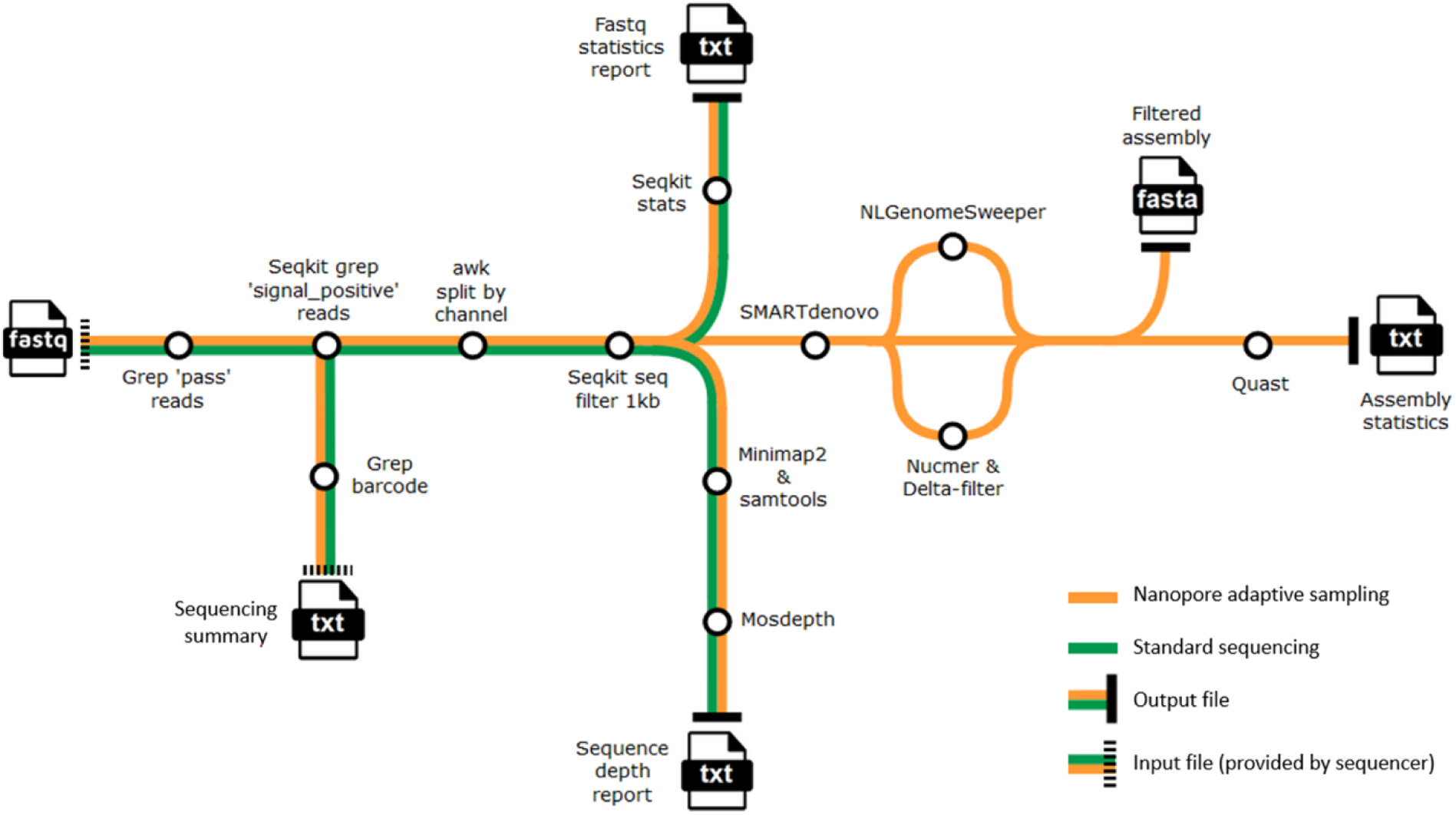
Workflow diagram summarizing the different steps involved in data processing and target regions assembly.

## Results

### ANSO77 AND DOUBLON *DE NOVO* GENOME ASSEMBLIES AND ANNOTATION

Metrics of the different steps during the assembly process are detailed in Additional files: Supplementary Data 1. Key metrics of the final assemblies are summarized in Table 1.

**Table 1.**
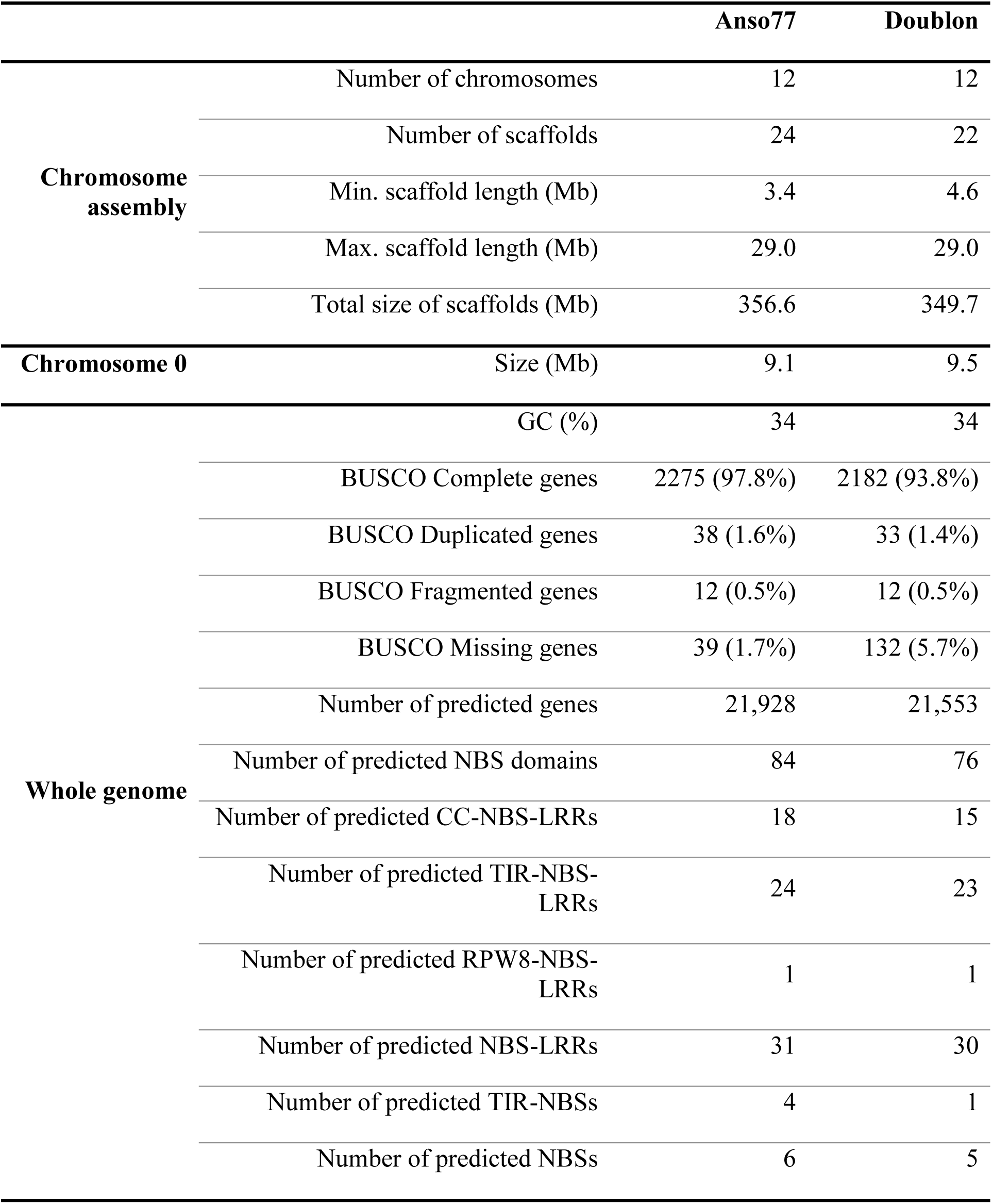
Summary metrics of the Anso77 and Doublon hybrid genome assemblies and annotations.

We annotated 21,928 genes in the Anso77 genome, and we identified a functional annotation in 20,946 (95.5%) of them. In the Doublon assembly, we annotated 21,553 genes, with 20,561 (95.4 %) being functionally annotated. Genes had an average length of 4,297 and 4,265 bp, with 6.09 and 6.03 exons in average per gene in Anso77 and Doublon, respectively.

We predicted 84 and 76 NBS domains in the genomes of Anso77 and Doublon, respectively. These numbers fall within the values obtained in other previously published melon genomes (Additional file 1: Table S1). Based on InterProScan (Hunter et al., 2009) domain identification in the 10 kb flanking sequence on both sides of the NBS domain, potential genes containing the predicted NBS domains were classified into different categories (Table 1).

The accuracy of the assemblies concerning NLR genes was assessed by inspecting the accuracy of the *Vat* homologs, whose cDNA sequence was previously obtained by Sanger sequencing (Chovelon et al., 2021). For Anso77, the homologs *AN-Vat2*, *AN-Vat3* and *AN-Vat5* fully matched the assembly here generated, while *AN-Vat1* and *AN-Vat4* contained one SNP each. For Doublon, all the three *Vat* homologs fully matched the assembled sequence.

### NAS TARGET REGIONS CONSTRUCTION

We arranged the 84 NLR predicted domains on Anso77 into 15 groups comprising nine ROIs with 2-28 NBS domains and six ROIs with isolated NBS domains. We also found 15 groups and similar physical positions of the NLR genes on Doublon and on the previously published melon genomes. After adding the 20 kb flanking zones, the 15 target regions varied in sizes from ∼41 to ∼1,378 kb, representing a total length of ∼6.16 Mb of the ∼370 Mb Anso77 genome (∼1.68%) (Table 2).

**Table 2.** Detailed information about the 15 target regions of Anso77. The *Vat* region is included within region 08.

| Target region | Chr | Start position | End position | Size | Predicted NBS domains |
| --- | --- | --- | --- | --- | --- |
| Region 01 | 01 | 32,471,836 | 32,684,765 | 212,929 | 14 |
| Region 02 | 02 | 979,011 | 1,026,074 | 47,063 | 2 |
| Region 03 | 02 | 15,072,686 | 15,113,435 | 40,749 | 1 |
| Region 04 | 04 | 5,452,467 | 5,493,242 | 40,775 | 1 |
| Region 05 | 04 | 26,516,353 | 27,894,199 | 1,377,846 | 7 |
| Region 06 | 05 | 14,002,942 | 14,043,763 | 40,821 | 1 |
| Region 07 | 05 | 17,364,466 | 18,340,871 | 976,405 | 2 |
| Region 08 | 05 | 25,108,836 | 26,146,869 | 1,038,033 | 28 |
| Region 09 | 06 | 5,837,171 | 5,877,998 | 40,827 | 1 |
| Region 10 | 07 | 2,550,415 | 2,591,165 | 40,750 | 1 |
| Region 11 | 07 | 24,326,063 | 24,411,978 | 85,915 | 4 |
| Region 12 | 08 | 3,465,204 | 3,505,984 | 40,780 | 1 |
| Region 13 | 09 | 645,676 | 815,779 | 170,103 | 10 |
| Region 14 | 09 | 6,558,754 | 7,597,791 | 1,039,037 | 4 |
| Region 15 | 11 | 6,635,285 | 7,601,705 | 966,420 | 7 |

After masking REs, the final file provided to the PromethION sequencer comprised 935 target regions, ranging from 502 pb to 67.609 kb (Additional file 2: Supplementary Data). They accounted for a total of ∼5.23 Mb of the ∼370 Mb Anso77 genome (∼1.41%).

### NAS TARGET REGIONS VALIDATION: EFFECTIVE ENRICHMENT OF NLR CLUSTERS IN MELON

Over 8.62 million reads and 19.86 Gb (42.71%) belonged to Anso77, and over 11.97 million reads and 26.63 Gb (57.27%) belonged to Doublon after barcodes trimming. We compared NAS to WGS of Anso77 and Doublon in terms of general metrics. For Anso77, further read splitting by channel and filtering of this data by quality, “end reason” and length resulted in 110.06 K reads resulting on 1.14 Gb coming from the NAS half-flowcell. The other half-flowcell (WGS) yielded 1.12 million reads with a cumulative size of 11.93 Gb for Anso77. Using the same processing for Doublon, 163.84 K reads generating 1.56 Gb from were assigned to the NAS half-flowcell, while 15.81 Gb from 1.70 million reads belonged to the WGS part. For both Anso77 and Doublon, the N50 value from the filtered NAS reads was very similar to the filtered WGS ones. All information about the generated datasets is grouped in Table 3.

**Table 3.** Anso77, Doublon and Chang-Bougi sequencing metrics. Anso77 and Doublon were sequenced using both NAS and WGS, while Chang-Bougi was only sequenced using NAS. Total WGS “pass” and total NAS “pass” represent the reads having a “PASS” flag (quality over 10) and assigned to the WGS and NAS half-flowcell, respectively. Filtered WGS and filtered NAS represent the two sets of reads just mentioned after filters of “end reason” and length.

| Dataset | Cultivar | Sequences number | Size (bp) | Mean length (bp) | Max. length (bp) | N50 | Q20 (%) | Q30 (%) |
| --- | --- | --- | --- | --- | --- | --- | --- | --- |
| Total dataset | Anso77 | 8,628,783 | 19,860,863,026 | 2,302 | 174,519 | 13,263 | 82.17 | 72.04 |
|  | Doublon | 11,976,866 | 26,639,849,880 | 2,224 | 193,630 | 11,968 | 83.65 | 73.58 |
|  | Chang-Bougi | 3,320,000 | 3,454,320,986 | 1,041 | 467,827 | 883 | 76.23 | 58.39 |
| Total WGS “pass” | Anso77 | 1,423,908 | 13,579,603,804 | 9,536 | 143,366 | 16,713 | 87.55 | 77.18 |
|  | Doublon | 2,033,986 | 18,285,705,305 | 8,990 | 137,509 | 15,261 | 87.55 | 77.33 |
| Total NAS “pass” | Anso77 | 6,634,798 | 4,807,120,593 | 725 | 126,590 | 617 | 87.89 | 77.41 |
|  | Doublon | 9,407,579 | 6,779,872,639 | 721 | 112,867 | 614 | 88.01 | 77.78 |
|  | Chang-Bougi | 3,056,000 | 3,115,590,337 | 1,020 | 232,918 | 878 | 82.81 | 64.01 |
| Filtered WGS | Anso77 | 1,122,605 | 11,939,444,337 | 10,636 | 143,366 | 16,772 | 88.00 | 77.75 |
|  | Doublon | 1,704,828 | 15,807,284,512 | 9,272 | 137,509 | 15,120 | 88.08 | 78.00 |
| Filtered NAS | Anso77 | 110,061 | 1,147,896,611 | 10,430 | 123,373 | 16,911 | 88.17 | 78.03 |
|  | Doublon | 163,843 | 1,561,078,862 | 9,528 | 112,867 | 15,182 | 88.09 | 77.99 |
|  | Chang-Bougi | 96,626 | 803,580,422 | 8,316 | 122,918 | 13,749 | 81.40 | 66.60 |

The length distribution of NAS-generated reads peaked at around 500 bp corresponding to rejected reads by adaptive sampling (91.20% and 92.20% of the total “pass” reads for Anso77 and Doublon) (Figure 3A). When rejected reads were out-filtered, a similar length distribution profile was observed for reads obtained in WGS and in NAS for both cultivars (Figure 3B, C).

**Figure 3.**
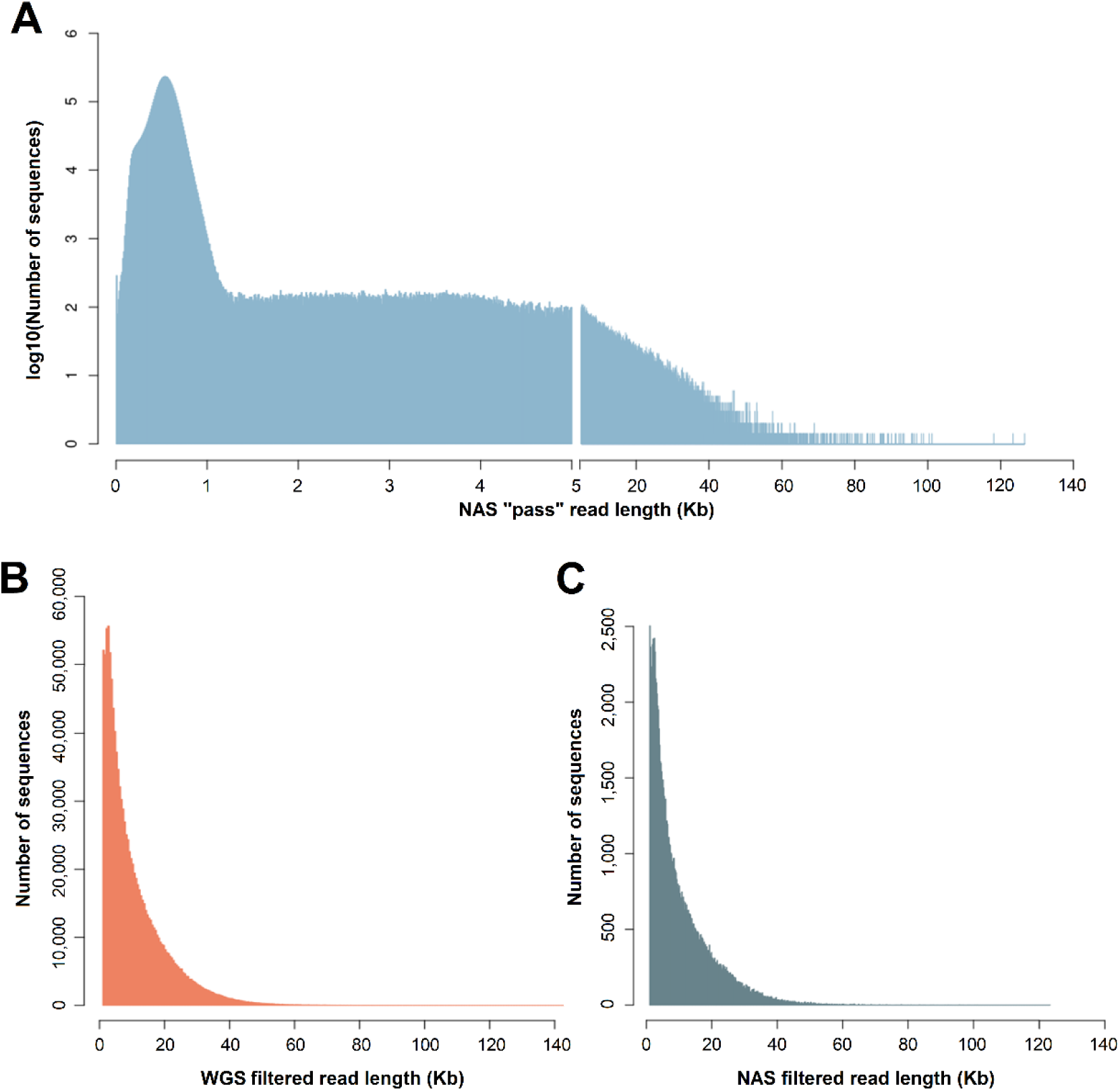
A) Length distribution of “PASS”-tagged NAS reads. The number of reads was log-transformed. B) Length distribution of WGS reads after filtering by “end reason”, quality and length. C) Length distribution of NAS reads after filtering by “end reason”, quality and length.

Focusing on Anso77, we evaluated the read depth on the target regions and on the rest of the chromosome in the NAS approach. The sequence depth on the target regions at the end of the experience was vastly higher than on the rest of the chromosome (Figure 4), with an average frequency of the target regions of 63.37. Sequence depth remained stable throughout the entire ROIs, even more so when the ROIs had a smaller size (Figure 4A-C). The standard deviation of the sequence depth on the ROIs ranged from 1.38 for region 03 to 20.19 for region 07. It presented values of 10.29, 16.15 and 1.80 for regions 01, 08 and 12, respectively. Actually, the increase of sequence depth was gradual on the 20 kb flanking, obtaining the highest depth on the ROIs. This increase exhibited a similar behavior regardless of the ROI’s size (Figure 4D).

**Figure 4.**
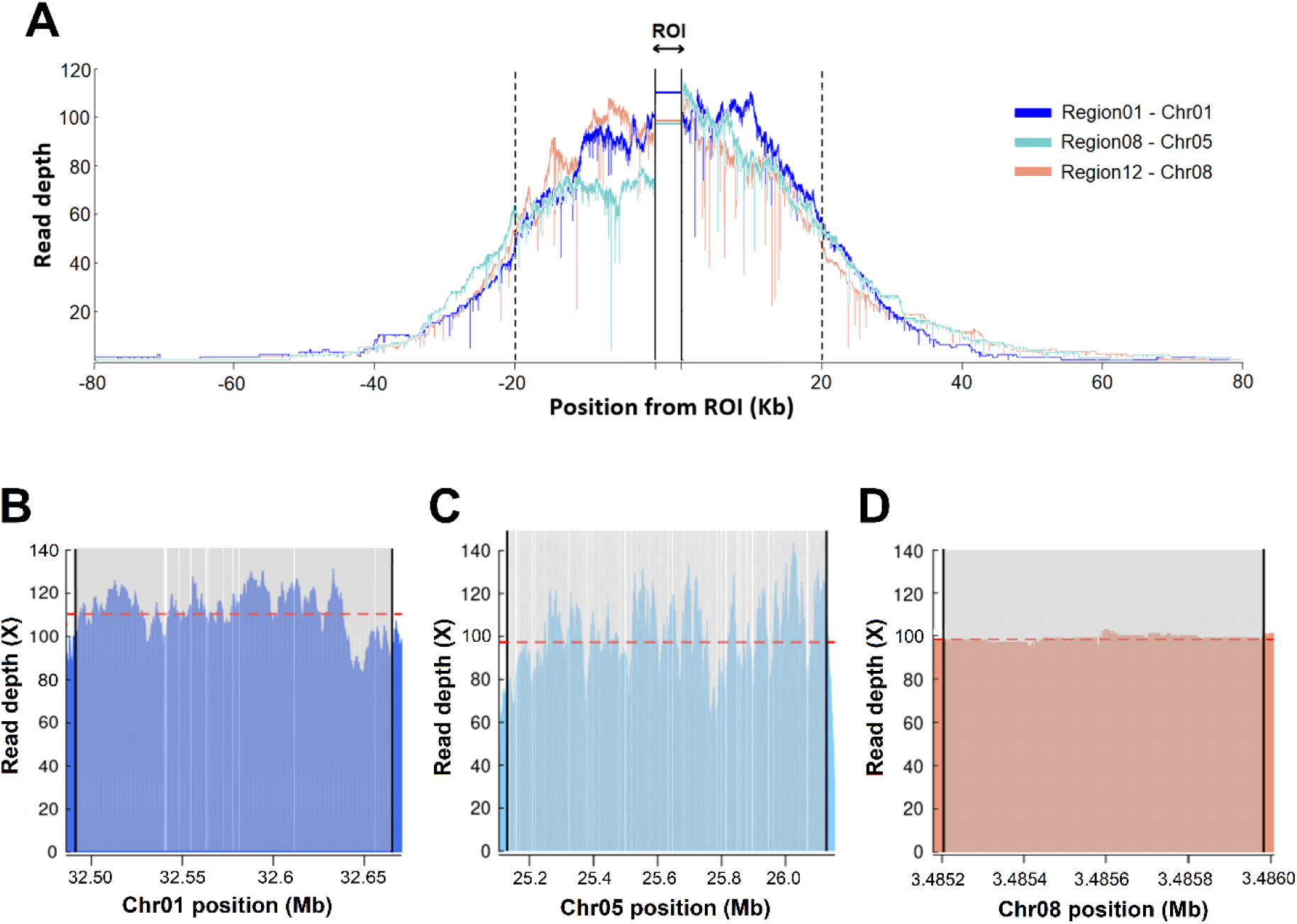
A) NAS sequencing depth on three target regions of different sizes compared to the rest of the chromosome. Target regions (ROI + 20kb buffer) are represented between black dotted bars, while ROIs are collapsed and represented between black solid bars. B) NAS sequencing depth on the ROI of region 01 (≈173 kb). C) NAS sequencing depth on the ROI of region 08 (≈998 kb) D) NAS sequencing depth on the ROI of region 12 (≈1 kb). Region 08 contains the well-studied *Vat* cluster. For B, C and D, vertical colored bars represent the enriched regions, while vertical white bars represent masked repetitive elements.

The half-flowcell design allowed us to calculate the enrichment obtained in NAS compared to a WGS approach. We obtained an enrichment by yield for Anso77 variable among target regions, ranging from 2.45 to 5.18 at the end of the run (Figure 5A). Considering the enrichment by selection, we observed an increase ranging from 45.56 to 102.91 (Figure 5B). On average, we obtained an enrichment by yield and an enrichment by selection of 3.96 and 78.38, respectively (Table 4). Target regions were sequenced at a lower rate than the rest of the genome in WGS, with a relative frequency of 0.81 (Table 4). Regarding enrichment over time, we observed that both the enrichment by yield and enrichment by selection followed very similar patterns. This enrichment reached its maximum at the beginning of the run for most of the regions when most of the flowcell channels were actively sequencing, and it decreased over time. This fact reflects that channel inactivation occurred faster on the NAS half-flowcell.

**Figure 5.**
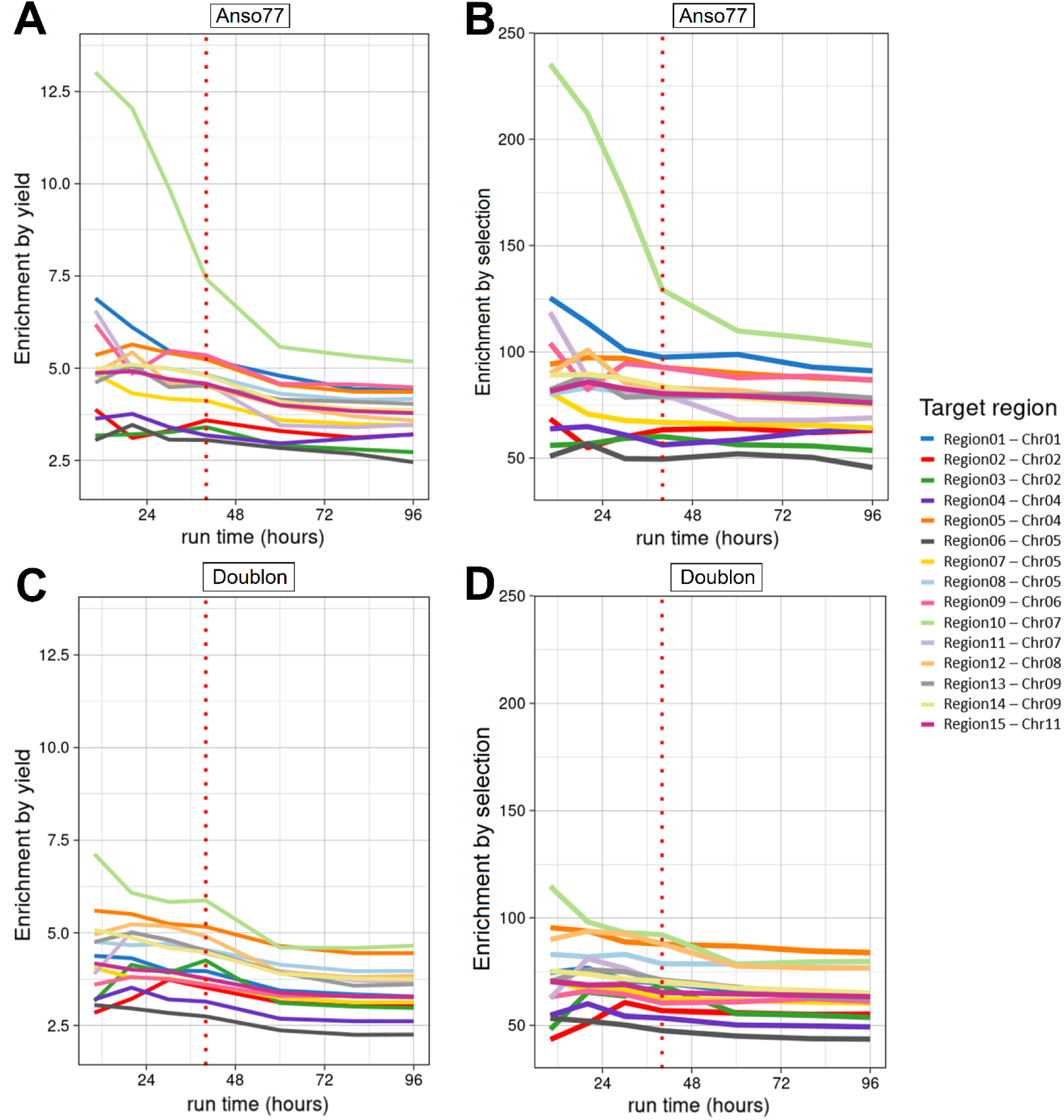
Enrichment by yield (A, C) and enrichment by selection (B, D) of the 15 target regions from Anso77 (A, B) and Doublon (C, D). Vertical red-dotted bars denote the flowcell washing flush time.

**Table 4.** Summary statistics of the NAS and WGS runs for Anso77 and Doublon. Only the chromosomes including target regions were considered for calculating the average off-target depth.

|  | **WGS**<br>**Anso77** | **NAS**<br>**Anso77** | **WGS**<br>**Doublon** | **NAS**<br>**Doublon** |
| --- | --- | --- | --- | --- |
| **Average on-target depth** | 22.79 | 90.18 | 31.74 | 118.28 |
| **Average off-target depth** | 28.19 | 1.42 | 32.52 | 1.73 |
| **Relative frequency of target regions** | 0.81 | 63.37 | 0.98 | 68.21 |
| **Average enrichment by yield** |  | 3.96 |  | 3.73 |
| **Average enrichment by selection** |  | 78.38 |  | 69.92 |

Region 10 exhibited a very particular behavior, being extremely enriched at the beginning of the run (Figure 5A, B). As shown in Additional files: Figure S1A, this high enrichment during the first hours of the run corresponded to a poor sequencing in the WGS approach. Furthermore, the NAS approach provided in just 10 hours a sequence depth comparable to that achieved in the entire WGS run (Additional files: Figure S1A). The washing flush contributed to an increase in pore activity in both NAS and WGS approaches (Additional files: Figure S1A).

To confirm the applicability of NAS in targeting the entire spectrum of NLR clusters in melon, we extended its use to Doublon, a cultivar within the same subspecies as Anso77 but belonging to a distinct botanical group. Similar to Anso77, the NAS sequence depth on the target regions at the end of the run always exceeded (2.25 to 4.65X) that of the WGS approach (Figure 5C, D). The least (region 06) and most (region 10) enriched regions were the same for both cultivars. Notably, the enrichment by yield presented a Kendall’s coefficient of concordance (W) of 0.87 between Anso77 and Doublon (p-value of 0.04), suggesting region-specific patterns rather than cultivar-based differences.

Target regions of Doublon were sequenced and mapped with an identical ratio (0.98) to the rest of the genome in WGS (Table 4). Regarding the enrichment by selection in Doublon, we obtained values ranging from 43.52 to 83.89 fold, maintaining a significant correlation with the values previously observed for Anso77 (W=0.89; p-value=0.04). Overall, we demonstrated an average enrichment by yield between all the target regions of 3.73 and an average enrichment by selection of 69.92 (Table 4). In terms of enrichment over time, the results were consistent with those obtained for Anso77, encompassing both the enrichment by yield and enrichment by selection (Figure 5C,D).

### ONT ADAPTIVE SAMPLING ALLOWS THE CORRECT ASSEMBLY OF NLR CLUSTERS IN ANSO77 AND DOUBLON

The assembly of NAS-enriched reads provided very contiguous and accurate assemblies of the target regions in both Anso77 and Doublon. Each cultivars presented a single contig for each target region. NAS assembly metrics are grouped in Table 5. Notably, the size of all contigs was over the size of their corresponding target region.

**Table 5.**
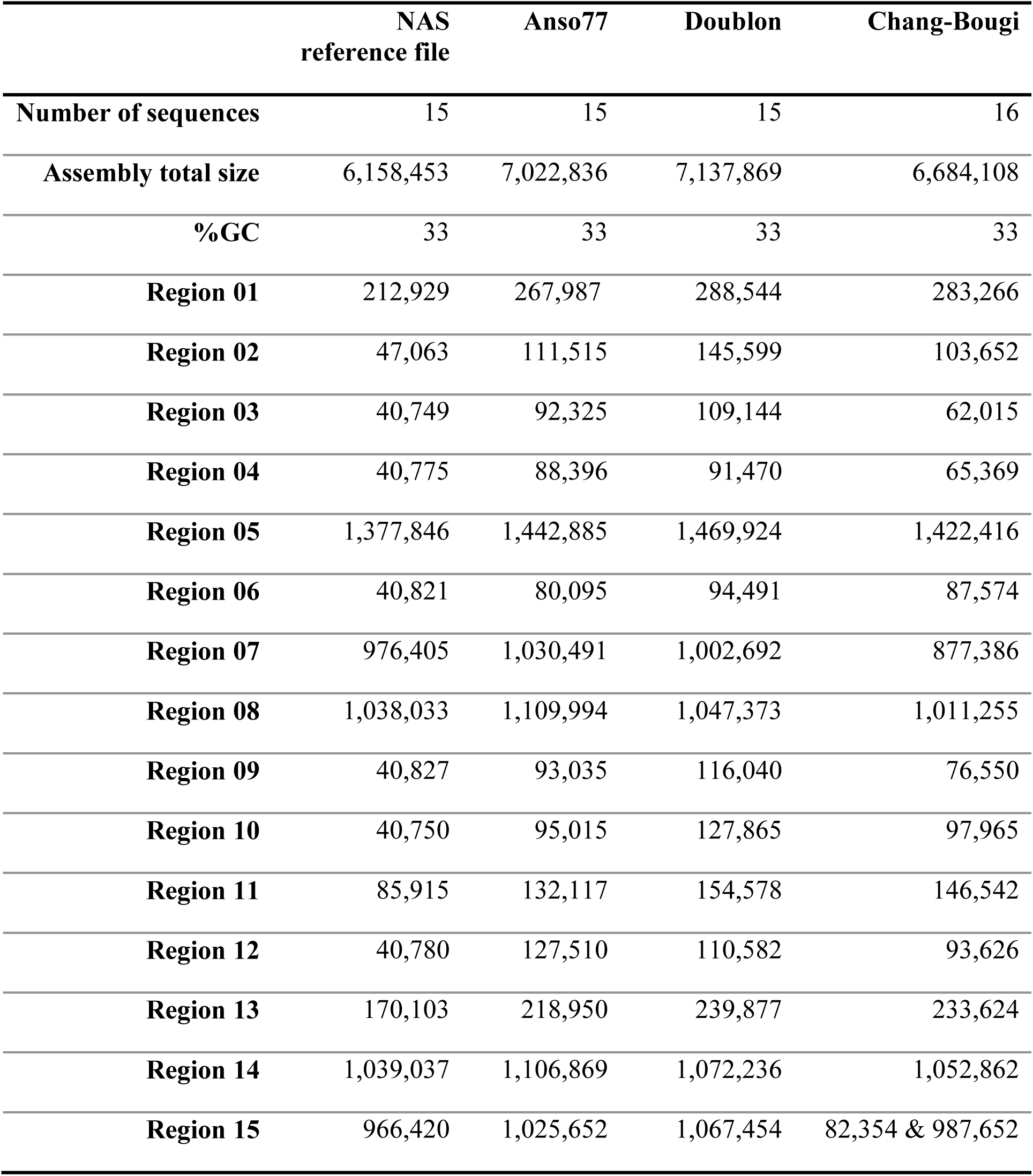
Anso77 and Doublon NAS assembly metrics. Regions of the NAS reference file represent the size (bp) of the target regions provided to the sequencer. Regions of Anso77, Doublon and Chang-Bougi represent the size (bp) of the contigs matching those regions.

|  | NAS<br>reference file | Anso77 | Doublon | Chang-Bougi |
| --- | --- | --- | --- | --- |
| **Number of sequences** | 15 | 15 | 15 | 16 |
| **Assembly total size** | 6,158,453 | 7,022,836 | 7,137,869 | 6,684,108 |
| **%GC** | 33 | 33 | 33 | 33 |
| **Region 01** | 212,929 | 267,987 | 288,544 | 283,266 |
| **Region 02** | 47,063 | 111,515 | 145,599 | 103,652 |
| **Region 03** | 40,749 | 92,325 | 109,144 | 62,015 |
| **Region 04** | 40,775 | 88,396 | 91,470 | 65,369 |
| **Region 05** | 1,377,846 | 1,442,885 | 1,469,924 | 1,422,416 |
| **Region 06** | 40,821 | 80,095 | 94,491 | 87,574 |
| **Region 07** | 976,405 | 1,030,491 | 1,002,692 | 877,386 |
| **Region 08** | 1,038,033 | 1,109,994 | 1,047,373 | 1,011,255 |
| **Region 09** | 40,827 | 93,035 | 116,040 | 76,550 |
| **Region 10** | 40,750 | 95,015 | 127,865 | 97,965 |
| **Region 11** | 85,915 | 132,117 | 154,578 | 146,542 |
| **Region 12** | 40,780 | 127,510 | 110,582 | 93,626 |
| **Region 13** | 170,103 | 218,950 | 239,877 | 233,624 |
| **Region 14** | 1,039,037 | 1,106,869 | 1,072,236 | 1,052,862 |
| **Region 15** | 966,420 | 1,025,652 | 1,067,454 | 82,354 & 987,652 |

We generated dot plots for Anso77 and Doublon comparing each assembled contig with the corresponding region in the whole genomes assembly we produced (Additional files: Figure S2). A perfect collinearity appeared in the dot plot of each target region, showing the fidelity of the NAS assemblies to the reference genomes. Notably, the same number of NBS domains and at the same positions were predicted in the NAS assembly compared to the reference genome for both cultivars, emphasizing the reliability of the NAS assemblies.

To further assess the accuracy of the NAS assemblies, we focused our attention on the well-known *Vat* region. Dot plots representing the *Vat* regions are shown in Figure 6. The dot plots heightened the complexity of this area with numerous duplicated sequences, but a perfect diagonal appeared between the reference and the NAS-assembled sequences. Moreover, we checked the sequence of the *Vat* homologs previously sequenced by Sanger sequencing (cDNA sequencing). Among the five homologs of Anso77, *AN-Vat1, AN-Vat2*, *AN-Vat3* and *AN-Vat5* fully matched the assembly generated with the NAS library, while *AN-Vat4* contained one SNP in the first exon (G/T on position 402710). This overcomes the reference assembly here presented, which contained the SNP in *AN-Vat4* but also one SNP in *AN-Vat1.* For Doublon, the three *Vat* homologous presented 100% DNA sequence similarity with the NAS assembly. Altogether, we demonstrated that NAS produces very contiguous and accurate assemblies in highly complex clusters of resistance genes.

**Figure 6.**
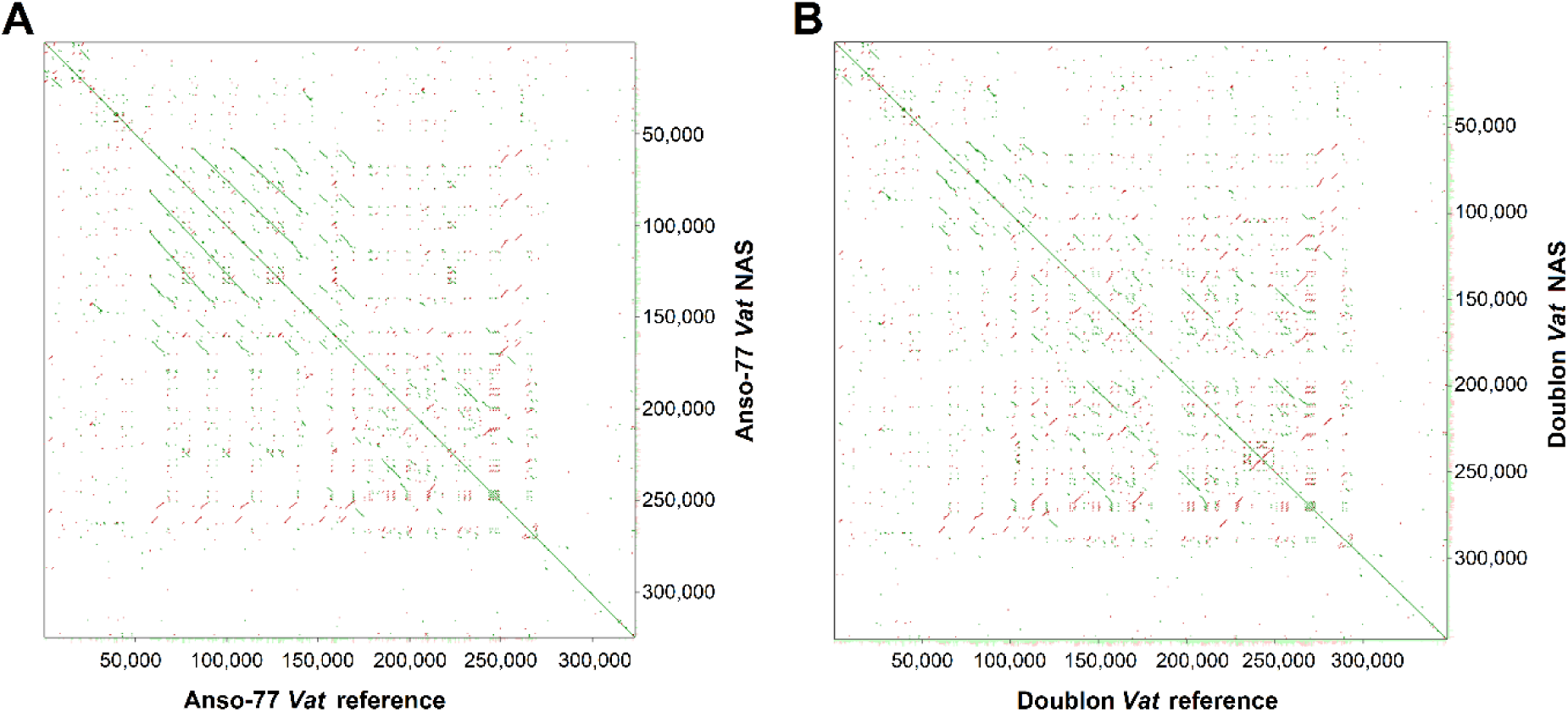
Dot plots representing the *Vat* region for Anso77 (A) and Doublon (B). Reference *Vat* region is represented on the x-axis, while the NAS-reconstructed *Vat* region is represented on the y-axis.

### TOWARDS AN ESTABLISHED PROCEDURE: ENRICHMENT AND ASSEMBLY OF NLR CLUSTERS FROM A DISTANT CULTIVAR PROVIDE VERY VALUABLE STRUCTURAL INFORMATION

We obtained 3.32 million reads and 3.45 Gb for Chang-Bougi in a 1/10 flowcell. After eliminating the reads rejected by adaptive sampling and filtering by quality and 1 kb length, 96.62 K target reads and 0.80 Gb were available for further processing. These reads exhibited an N50 of 13.75 kb, a measure comparable to that obtained for Anso77 and Doublon. Rejected reads had an average length of ∼790.52 bp, longer than that obtained for Anso77 and Doublon, due to the updated sequencing speed (400 bp/s).

The sequence depth on the target regions, mapped on the assembled contigs, averaged 41.82X. As illustrated in Figure 7, this depth was variable between regions, with the highest depth in region 05 (50.37X) and the lowest in region 03 (21.08X). However, the obtained sequenced depth between regions kept a significant concordance with that obtained for Anso77 and Doublon (Additional files: Figure S1-left) (W=0.78; p=0.003).

**Figure 7.**
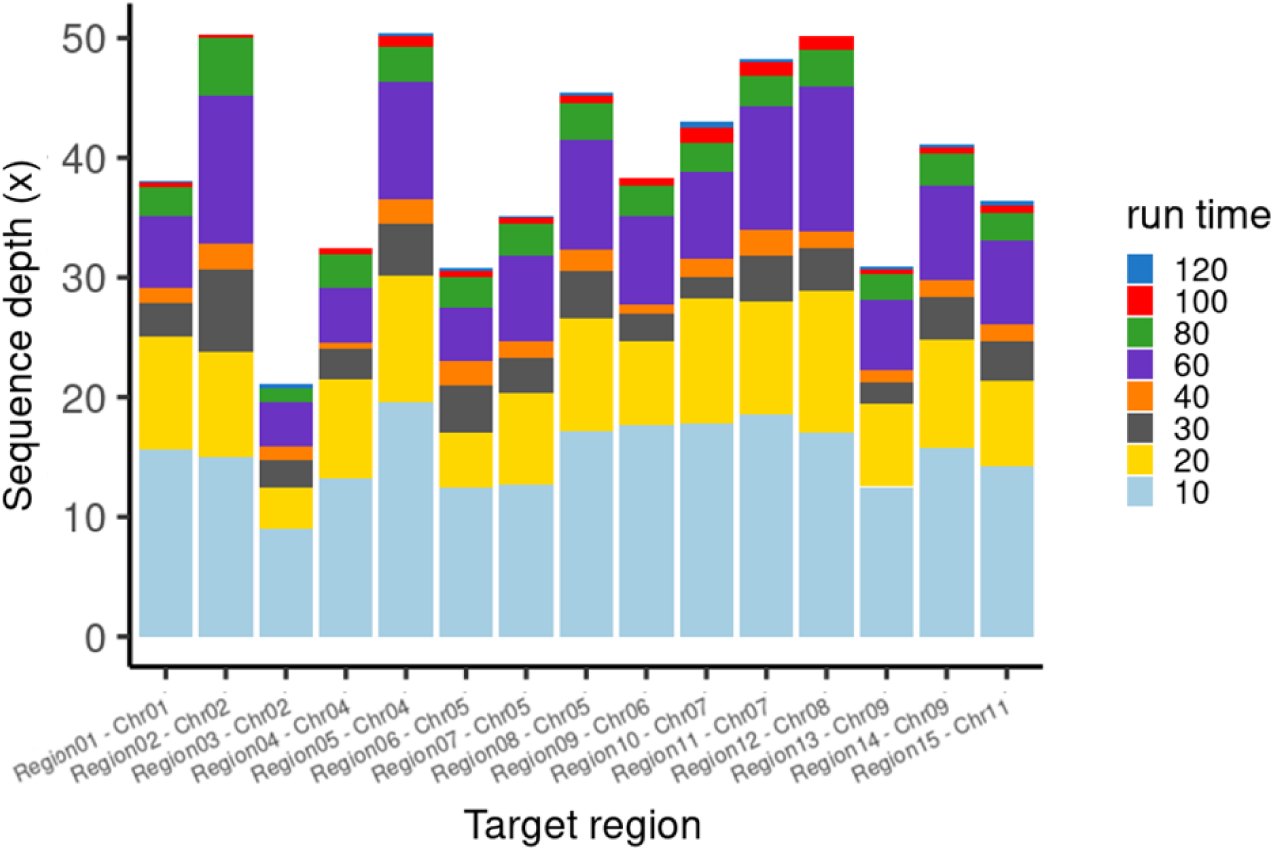
Sequence depth in the NAS experience of the 15 target regions from Chang-Bougi.

For Chang-Bougi, the assembly of NAS-enriched reads with SMARTdenovo resulted in 17 contigs. Notably, we observed an inversion of ≈100 kb on region 01 compared to Anso77 (Figure 8A). Region 13 and region 15 were fragmented into two contigs. Among the tested assemblers, Canu achieved a contiguous assembly of region 13 in a single contig (Additional files: Figure S3). Consequently, we retained region 13 from the Canu assembly. No assembler succeeded in reconstructing region 15 into a single contig. We then investigated why region 15 was fragmented into two contigs. Due to the high degree of fragmentation of the Illumina-based published assembly, we could not conclude after contig alignment. As Chang-Bougi belongs to the *makuwa* botanical group, we mapped the two contigs to the publicly available genomes from this group: Early Silver Line, Ohgon and Sakata’s Sweet (Oren et al., 2022). We identified a very large and size-conserved insertion (ranging from 862 to 871 kb) at the breakpoint between the two contigs obtained for Chang-Bougi (Figure 8A). No NBS domain was predicted in this insertion on the genomes of Early Silver Line, Ohgon and Sakata’s Sweet. We could not recover this insertion in Chang-Bougi as it was not present in the provided reference. The total size of the assembly was 6.68 Mb. Table 4 shows the detailed NAS assembly metrics.

**Figure 8.**
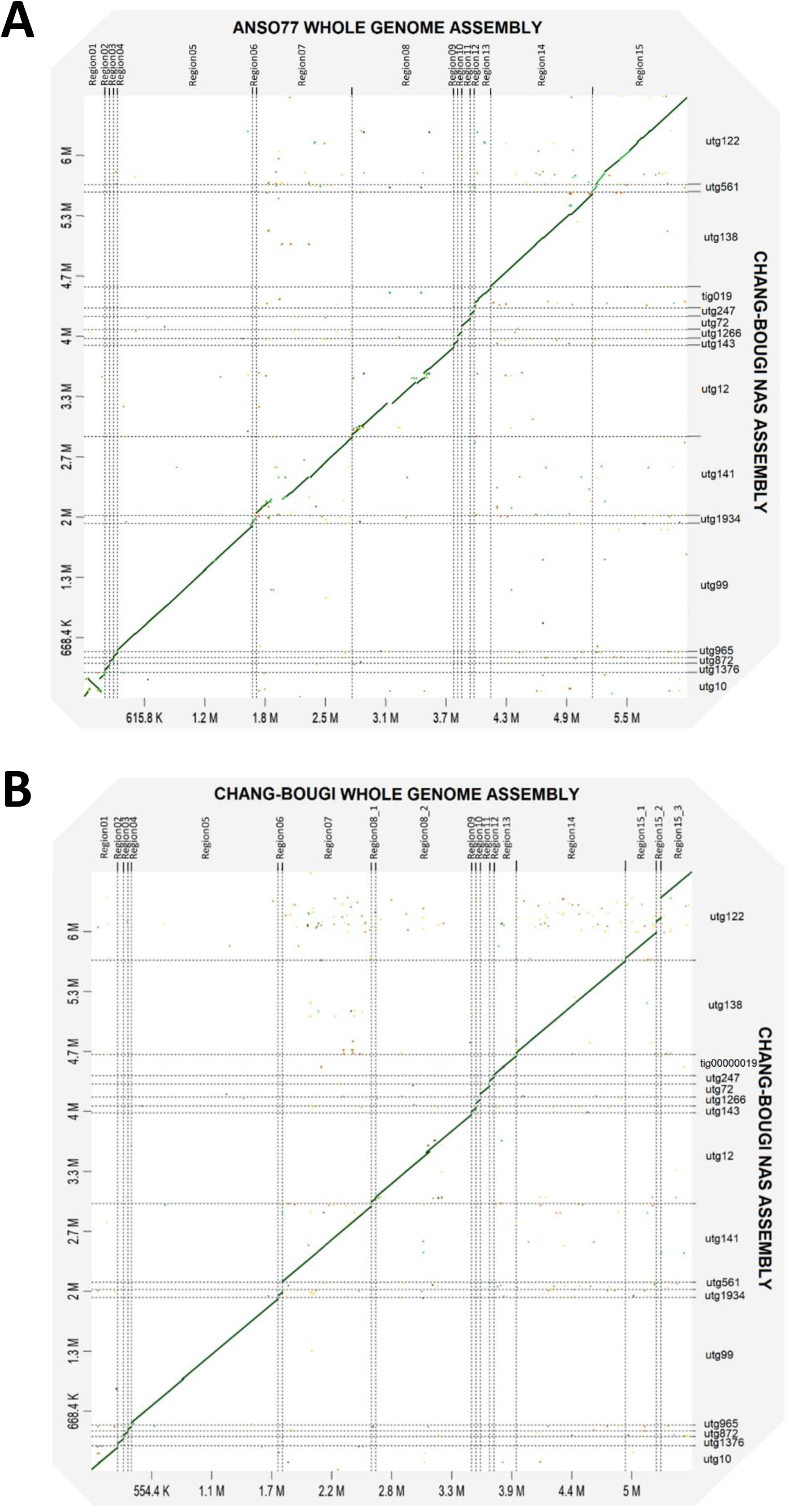
Dot plots representing the NAS filtered assembly of Chang-Bougi (y-axis) against the 15 target regions from Anso77 (A) and the 18 NLR clusters from Chang-Bougi (B).

In the draft genome of Chang-Bougi generated by (Shin et al., 2019), we identified 81 NBS domains spread across 18 contigs. These contigs matched the previously identified 15 ROIs of Anso77 with no extra clusters (Figure 8B). In the NAS assembly, we identified 83 NBS domains. The two extra NBS domains predicted in the NAS assembly were located in region 08, one in the *Vat* region and the other outside. We manually annotated the *Vat* region of both Chang-Bougi assemblies (NAS and published) revealing some discrepancies in the complex and repetitive area between *Vat1* and *VatRev* (Figure 9A). In the NAS assembly, we identified the extra NBS domain within the *Vat* region as a *Vat* homolog with four R65aa motifs (Figure 9A). We confirmed the presence and the structure of this *Vat* gene with four R65aa motifs as well as the presence of the *Vat* genes with three R65aa through PCR using the published primers Z649FR and Z1431FR (Figure 9B). Finally, the presence of long reads encompassing the pairs of genes *Vat1*:*Vat2* and *Vat2*:*Vat3* confirmed the NAS-assembled structure.

**Figure 9.**
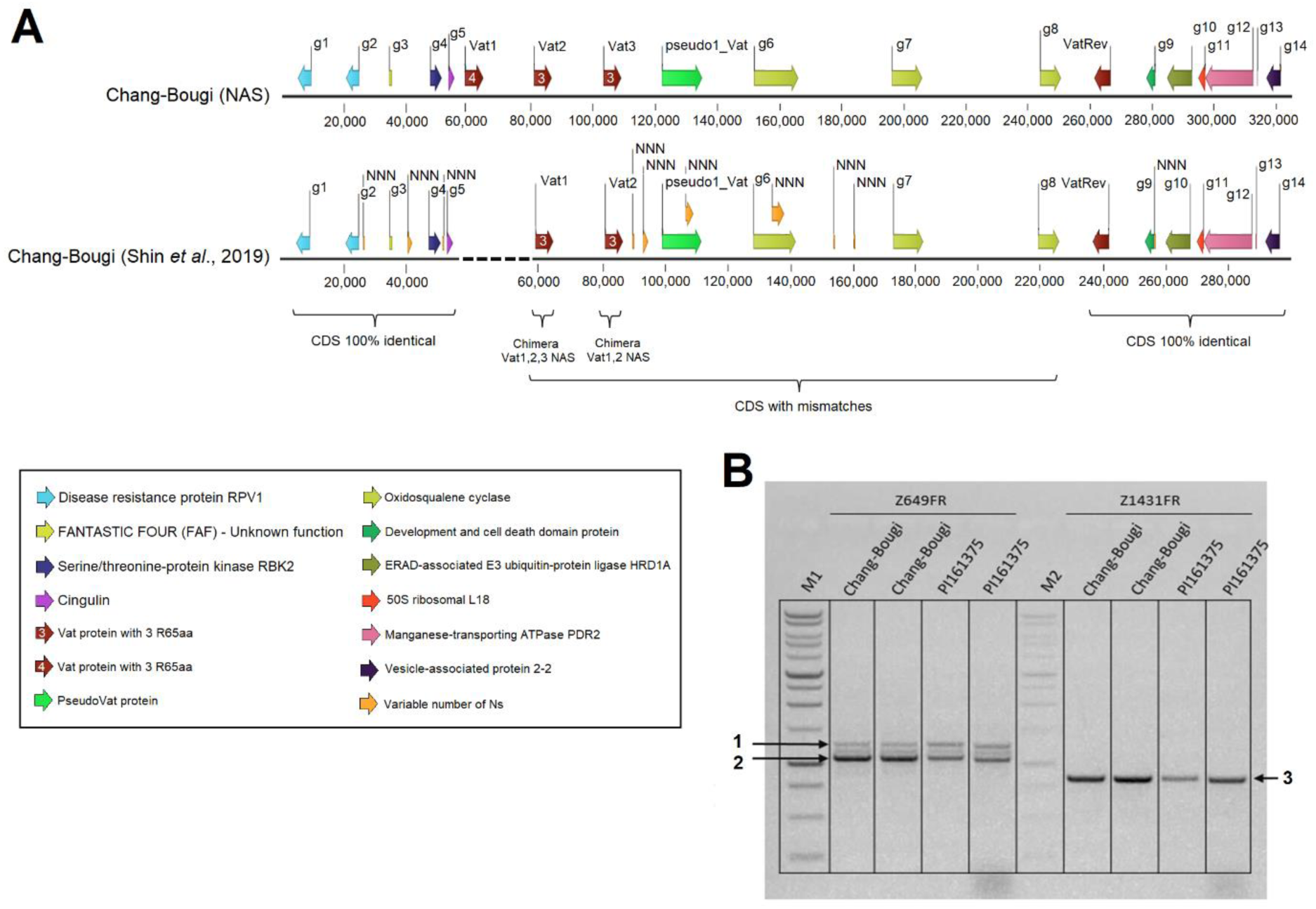
A) Genes identified after manual annotation within the *Vat* regions of Chang-Bougi. The sequence above was obtained from the NAS assembly, while the sequence below was recovered from the publicly available genome assembly (Shin *et al*., 2019). B) Agarose gel electrophoresis of PCR products obtained using primers Z649FR and Z1431FR. Lanes M1 and M2 are the two 1 kb DNA ladders (Promega, Madison, WI, USA). PI161375 was used as a control having a *Vat1* with four R65aa motifs and a *Vat2* with three R65aa motifs. Bands pointed with arrows represent an amplicon of four R65aa motifs (1), an amplicon of three R65aa motifs (2), and a specific amplicon of four R65aa motifs (3).

## Discussion

Herein, we highlighted NAS as a promising approach for studying the polymorphisms of complex genomic ROIs. We chose melon as a model for which we selected highly diverse ROIs in terms of size and NLR gene content.

Several factors can influence the efficiency of NAS. Among them, we set up two key considerations before launching the NAS experiments. First, we hypothesized a direct relationship between the ideal size of sequenced fragments and the size of the ROIs (Figure 10). Given that ROIs sizes varied here from ≈41 to ≈1378 kb, we used standard DNA extractions (10-30 kb) as they were expected to produce a more stable sequencing depth on ROIs than ultra-long reads (100-300 kb) for the same yield. We believe that this approach reduces off-target sequencing and avoids channels blockage when rejecting very long reads, as established in ONT recommendations (Community Nanopore, 2023). Second, REs cover an important portion of the melon genome (Castanera et al., 2020), and they are especially frequent inside the NLR gene clusters (Chovelon et al., 2021). To prevent sequencing off-target REs with high sequence similarity to those within the initial target regions, we assumed that masking repetitive elements within the provided target regions would diminish the quantity of off-target data, contributing to increased enrichment. Among the 6.16 Mb, we masked 0.93 Mb of repetitive sequences. Masking REs in genomes prior to NAS was previously suggested (Zhang et al., 2021).

**Figure 10.**
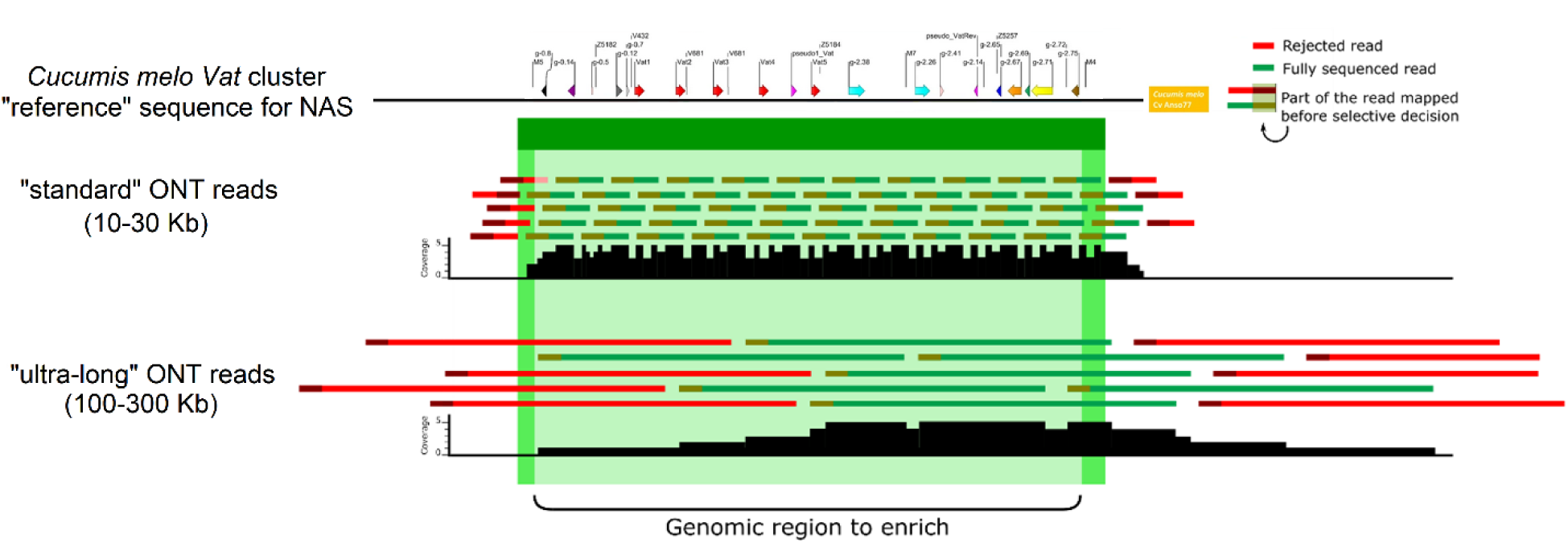
Diagram illustrating the difference in coverage and extent of the area outside the region to be enriched for standard size fragments (10-30kb) and for ultra-high molecular weight fragments (100-300kb) on the *Vat* (melon) cluster. For the same yield (illustrated here by an arbitrary overall depth of 5X), standard fragments make it possible to achieve a depth more concentrated on the area to be enriched and to sequence less outside this area. For convenience, only reads oriented from 5’ to 3’ are represented.

We investigated the interest of NAS compared to WGS. One key factor that largely affects final yield in ONT sequencing runs is the number of active channels at the beginning of the run and their lifespan, which may significantly differ from one flowcell to another. Therefore, we implemented a half-flowcell design to compare NAS and WGS eliminating biases that may arise when using two different flowcells, as previously settled in many studies (De Meulenaere et al., 2022; Martin et al., 2022). Results showed that for both Anso77 and Doublon, NAS produced about four times more on-target data than WGS (Table 3), while generating about ten times less total data (see filtered NAS and WGS in Table 2). Nevertheless, the sequence depth between target regions was more variable in NAS than in WGS, although this did not compromise the accurate assembly of the ROIs in all the cultivars. We found no correlation between target size and sequence depth (Additional files: Figure S4A) but there was a moderate correlation between percentage of masking and sequence depth (additional files: Figure S4B).

We proposed two measures of enrichment adapted from previous studies that have tested NAS on metagenomics samples or panels of many small regions of the genome (Hogers et al., 2020; Martin et al., 2022): the enrichment by yield, a widespread and simple metric; and the enrichment by selection, a metric not biased by the sequencing behavior of each target region. These enrichment measures make sense in our study because our goal is to increase coverage in the complex ROIs to generate more accurate assemblies. We obtained an enrichment by yield up to 3.7 times on average and an enrichment by selection up to 69 times on average, even when the reference was genetically distant from the sequenced accession. These findings are comparable to the best results previously obtained enriching individuals in metagenomics samples (De Meulenaere et al., 2022; Martin et al., 2022) and outperformed the enrichment values previously obtained with loci panels (Hogers et al., 2020). The successful enrichment with NAS could be linked to the percentage of the genome targeted here (∼1.41%), the size of the targets, or the size of the DNA fragments, as they have been demonstrated to be main factors of enrichment rate (Martin et al., 2022; Community Nanopore, 2023). In addition, a late nuclease flush performed when the percentage of sequencing pores was around 10% might have contributed to the good performance, instead of doing it at a fixed time (Payne et al., 2021; Martin et al., 2022; Nakamura et al., 2023). The enrichments over time of the different target regions were higher and more variable at the beginning of the run but globally stabilized after 70h (Figure 5). Actually, channel inactivation occurred faster on the NAS half-flowcell, either due to the repetitive potential flipping to reject off-target sequences or simply because the likelihood of channel clogging is statistically related to the number of sequenced molecules (Kovaka et al., 2021; Martin et al., 2022).

Previous studies have used NAS to enrich specific species in metagenomics samples (Kipp et al., 2021; Martin et al., 2022; Ulrich et al., 2022) and relatively small sequences within an organism, such as exon panels or panels of loci of key variants (Hogers et al., 2020; Filser et al., 2023; Nakamura et al., 2023). Here, we demonstrated the power of NAS as a tool for enriching ROIs that represent isolated NLR genes or complex clusters of NLRs in a plant crop species. The correct assembly of these complex regions typically requires long reads as those provided by ONT or dedicated laborious approaches such as the R gene enrichment sequencing (RenSeq) method (Witek et al., 2016; Van de Weyer et al., 2019; Huang et al., 2022; Vendelbo et al., 2022; Adams et al., 2023). In fact, NLR genes may have been miss-predicted, especially when short-read sequencing technologies were used. When deciphering the *Vat* region of DHL92 (Garcia-Mas et al., 2012), 2 to 4 functional NLR genes and some pseudo NRLs were found (Chovelon et al., 2021). After looking for these NLRs in the early released genome of DHL92, it appeared they were misassembled. Only when a high-quality genome (with long reads, optical maps or HiC) was released (Castanera et al., 2020), the *Vat* genes were finally congruent with Sanger sequencing from long-range PCR (Chovelon et al., 2021). Moreover, in the present study, we compared the *Vat* cluster in the cultivar Chang-Bougi derived from short WGS (Shin et al., 2019) and long NAS reads. We showed that the WGS assembly was erroneous in terms of the number and sequences of homologous genes (Figure 9) and that the NAS assembly accurately reconstructed the region.

NLR genes are encompassed within the dispensable portion of the genome (Barragan & Weigel, 2021; Shang et al., 2022), and therefore the NLR reference used for NAS should be carefully selected when targeting the NLRome of a species. Our prediction of the number of NBS domains in Anso77 and Doublon, along with all previously published melon genome assemblies, consistently yielded similar values (Additional files: Table S1). Across all cases, we did not find more than 15 groups of NLR genes regardless of the subspecies addressed, *melo* or *agrestis*. These findings indicate a well-conserved number and location of NLR genes in melon. We chose Anso77, a Spanish cultivar belonging to the subspecies *melo* and the botanical group *inodorus*, as the reference for the NAS approach because it contained the highest number of *Vat* homologs within the *Vat* region used for benchmarking (Chovelon et al., 2021). The results here obtained with Doublon and Chang-Bougi suggested that this strategy was judicious. Doublon is a French melon line, belonging to the subspecies *melo*, and the botanical group *cantalupensis*. Using NAS without any short-read polishing we obtained a *Vat* cluster almost identical to the one derived from an assembly using HW-DNA, PacBio and ONT long sequences, Illumina short sequences, and optical maps. Chang-Bougi is a Korean melon line, belonging to the subspecies *agrestis*, and the botanical group *makuwa*. The *Vat* cluster we obtained using NAS was highly consistent with the *Vat* cluster of PI 161375 (Chovelon et al., 2021), a Korean line belonging to *agrestis* subspecies. However, a limitation appeared for very large SVs not present in the reference, as exemplified by the one identified in the chromosome 11 of Chang-Bougi that turned out to be present in the oriental melon clade. Using different high-quality reference genomes or even combining them into an “artificial” reference genome could address this limitation. Existing software such as BOSS-RUNS (Weilguny et al., 2023) already enables dynamically updating the decision strategies during the run, thereby allowing a better balance in the depth of target regions or multiplexed samples. Nonetheless, it remains uncertain whether NAS would have been able to discover extra NLR gene clusters if they existed. This should be possible if the additional NBS domains are conserved enough to match the provided reference.

Altogether, our study provides a blueprint for the selective capture of the NLRome in melon and could be extended to other important crop species. NLR gene numbers are generally low in the Cucurbitaceae family (Baggs et al., 2017; Barragan & Weigel, 2021), and they represent an ideal percentage of the genome to be targeted using NAS. However, this ideal situation does not correspond to reality in other species, as the number of NLR genes is highly variable between plant species independently of their genome size (Barragan & Weigel, 2021). To adapt the NAS procedure implemented here for NLR-rich plant species, certain adjustments should be made to align with the ideal targeted percentage of the genome. First, a reduction in the length of flanking regions surrounding the ROIs is recommended. Subsequently, employing a more rigorous definition of NLR clusters would help to reduce the percentage of targeted genome. Finally, a focus with NAS could be strictly done on the clustered NLRs, recovering the isolated NLRs with the low-pass rejected reads by NAS.

## Conclusions

NAS offered a flexible, real-time enrichment of selected clusters of NLR genes while reducing costs compared to a WGS approach. Such target enrichment did not require any laborious or expensive library preparation nor probes design and synthesis unlike previously developed target sequencing methods. This is particularly advantageous for researchers who may not have access to special molecular biology techniques or seek to conduct in-field experiments. NAS only requires an ONT sequencing device (that can be the low-cost MinION device), a reference genome, and one or several ROIs. In addition, the fast enrichment observed here may be of crucial interest when time is a critical factor. Moreover, we evidenced the ability of NAS to reduce the high off-target volume of data produced by WGS, addressing the growing challenges of data management and storage in the field of bioinformatics and genomics. This methodology, validated here on three melon cultivars, holds promise for its application across a large number of accessions. This is particularly relevant for breeding purposes as it opens avenues for creating multi-resistant varieties by tapping into the NLRome diversity.

## Acknowledgements

We thank Isabelle Dufau (INRAE-CNRGV) for furnishing high-quality HMW DNA and optical maps for Anso77 and Doublon. Moreover, we are grateful to Valerie Barbe for advice concerning the manuscript.

## Ethics approval and consent to participate

Not applicable.

## Consent for publication

Not applicable.

## Competing interests

The authors declare that they have no competing interests.

## Funding

This work was partly funded by the French *Ministère de l’agriculture et de la souveraineté alimentaire* (*Vat*&Co project - CASDAR − 2017-2021) and the French National Research Institute for Agriculture, Food and Environment, INRAE. The doctoral position of Javier Belinchon-Moreno is co-funded by the INRAE BAP Department and the EUR Implanteus of Avignon University, France.

## Authors’ contributions

J.B.M. performed the sequencing experiments with NAS, conducted the bioinformatics and statistical analyses, and carried out the NAS assemblies. P.F.R., N.B. and D.H. conceived the study. J.L., V.C., A.C. designed and identified the ROIs for NAS. A.B. and I.L. generated the ONT, Illumina and 10x genomic data for the whole genome assemblies. W.M. generated Bionano data for the whole genome assemblies, and participated in the hybrid scaffolding. J.L. and R.F.L. performed the whole genome assemblies. S.E. provided expertise and bioinformatics support. C.C. provided expertise and experimental support. V.R.R. performed manual annotation of the *Vat* cluster and did the PCR experiments. J.B.M., P.F.R., N.B. and D.H. wrote the manuscript. J.B.M. and A.C. did the data submission. All authors read and approved the final manuscript.

## Notes

### Competing Interest Statement

The authors have declared no competing interest.

