## Supplementary Methods, Data, Figures and Tables for "Nanopore adaptive sampling to identify the NLR-gene family in melon (*Cucumis melo* L.)"

#### DNA EXTRACTION, QUANTIFICATION, LINKED READ SEQUENCING AND LONG-READ SEQUENCING

DNA extraction, quantification, 10X Genomics linked-read sequencing (of Anso77) and long-reads sequencing (of Anso77 and Doublon) were reported in the supplementary methods S1 of (Chovelon et al., 2021). Raw 10X Genomics (Pleasanton, CA, USA) for Anso77, PacBio (Pacific Biosciences, Menlo Park, CA, USA) for Doublon and ONT (Oxford Nanopore Technologies, Oxford, UK) data for both cultivars were available in the NCBI database under the Bioprojects numbers PRJNA662717 for Anso77 and PRJNA662721 for Doublon. In this study, we generated new genomic resources described below and we applied different assembly methods to improve the genome sequences.

#### ILLUMINA LIBRARY PREPARATION AND SEQUENCING

For both cultivars, an Illumina sequencing library was prepared starting from DNA extracted with the Qiagen DNAeasy Plant miniKit (Qiagen, Valencia, CA, US). Sequencing was conducted using an Illumina NovaSeq 6000 system (Illumina, San Diego, CA, USA) with 2 × 150 bp paired-end reads according to the manufacturer's protocol recommendations. A phred quality score cutoff of 20 (corresponding to a 99% base-call accuracy) was applied to the generated fragments. Overlapping paired-end reads were merged into one single contig represented by a consensus sequence, to avoid considering that fragment twice in depth statistics.

#### BIONANO OPTICAL MAPPING DETECTION AND DATA PROCESSING

HMW-DNA labeling and staining was carried out according to the Bionano Prep Direct Label and Stain (DLS) protocol (BioNano Genomics, San Diego, CA, USA). Briefly, HMW DNA labeling was performed by incubating 750 ng of previously extracted genomic DNA with 1× DLE-1 Enzyme (BioNano Genomics, San Diego, CA, USA) for 2 hours in the presence of 1× DL-Green (BioNano Genomics, San Diego, CA, USA) and 1× DLE-1 Buffer (BioNano Genomics, San Diego, CA, USA). After proteinase K digestion and DL-Green cleanup, the DNA backbone was stained by mixing the labeled DNA with DNA Stain solution (BioNano Genomics, San Diego, CA, USA) in presence of 1× Flow Buffer (BioNano Genomics, San Diego, CA, USA) and 1× DTT (BioNano Genomics, San Diego, CA, USA), and incubating it overnight at room temperature. The DLS DNA concentration was quantified with a Qubit dsDNA HS Assay Kit (Invitrogen, Carlsbad, CA, USA). Labelled and stained HMW-DNA was loaded into a Saphyr chip (BioNano Genomics, San Diego, CA, USA). Loading of the chip and running of the Bionano Genomics Saphyr System (BioNano Genomics, San Diego, CA, USA) were performed according to the Saphyr System User Guide (BioNano Genomics, San Diego, CA, USA). Data processing was achieved using the Bionano Genomics Access software (BioNano Genomics, San Diego, CA, USA).

#### *DE NOVO* GENOME ASSEMBLIES AND HYBRID SCAFFOLDING

For Anso77, we merged ONT reads coming from the PromethION, MinION and Flongle devices. For Doublon, we combined PacBio and ONT reads. Adapters were removed from the ONT raw reads using porechop v. 0.2.4 (https://github.com/rrwick/Porechop). We performed an initial contig assembly for both cultivars using Canu v. 2.1 (Koren et al., 2017) with genomeSize=380m, corOutCoverage=100 and minReadLength=1000 settings. For Anso77, we performed two rounds of polishing with ema v. 0.6.2 (Shajii et al., 2018) and Pilon v. 1.23 (Walker et al., 2014) using the 10X Genomics linked-reads. We produced two rounds of polishing afterwards for both cultivars using the paired-end 2x150 bp Illumina reads, also with ema v. 0.6.2 and Pilon v. 1.23.

We assembled the Bionano raw data to construct the cmap file with Bionano Solve v. 3.6.1 (Bionano Genomics, San Diego, CA, USA). We used the generated cmap for both scaffolding and correction of incorrectly assembled contigs with Bionano Solve. This allowed the construction of scaffolds.

#### CHROMOSOME ALLOCATION

We manually assigned and ordered the scaffolds on the melon linkage groups by mapping forward-reverse primers of 148 highly conserved SSR markers between melon and cucumber (D. Li et al., 2011) (Additional files: Table S2) to construct the chromosomes. Localization of SSR markers on the scaffolds and Harukei-3 genome (Yano et al., 2020) was determined using a custom-made script (available on request).

#### GENOMES ANNOTATION

For both Anso77 and Doublon genomes, we performed *Ab initio* structural gene annotation using the deep-learning approach implemented in Helixer (Stiehler et al., 2021; Holst et al., 2023), available at https://www.plabipd.de/helixer_main.html. We removed contigs shorter than 25kb, and selected the land plant lineage for lineage-specific annotations. In addition, we used eggnog-Mapper v. 2.1.12 (Cantalapiedra et al., 2021), combined with eggnog DB v5.0.2 (Huerta-Cepas et al., 2019), to annotate the functions, Gene Ontology (GO), and Kyoto Encyclopedia of Genes and Genomes (KEGG) items of protein-coding genes. We used default parameters except '-m diamond --dmnd_iterate no --tax_scope eukaryota'.

For both lines, candidate NLR gene prediction based on the identification of the complete NB-ARC domain was performed using NLGenomeSweeper (Toda et al., 2020).

#### QUALITY CONTROLS

BUSCO version 3.0.2 (Simão et al., 2015) was used to assess the completeness of the Anso and Doublon assemblies, using the eudicotyledons_odb10 dataset with 2326 BUSCOs.

To check the accuracy of the assemblies concerning NLR genes, we focused our attention on the well-known Vat cluster located in chromosome 5 (Chovelon et al., 2021). We compared the assembled sequences to the sequence of the *Vat* homologs previously obtained by Sanger sequencing (Chovelon et al., 2021).

### Supplementary Data

Metrics of the generated reads for both Anso77 (ONT) and Doublon (ONT and PacBio) are summarized in Additional files: Table S3.

Initial long-read assembly followed by a polishing step using the 10X Genomics Linked-reads (for Anso77) and a polishing step using the Illumina paired-end reads yielded 159 contigs for Anso77 (N50 of 8.9 Mb) and 186 contigs (N50 of 15.2 Mb) for Doublon (Additional files: Table S4).

We generated a total volume of Bionano data of 902 and 805 Gb for Anso77 and Doublon, respectively. From this data, molecules larger than 150 kb represented 480 Gb data for Anso77 and 480 Gb data for Doublon, corresponding to 359,954 molecules with N50 of 235.8 kb and an average label density of 15.7/100kbp for Anso77 and 338,440 molecules with N50 of 256.7 kb and an average label density of 16/100kbp for Doublon (Additional files: Table S3). Thirty-three optical maps with a N50 of 20.4 Mb for a total genome map length of 388.8 Mb were produced for Anso77 and 32 maps with a N50 of 20.6 Mb for a total genome map length of 391.3 Mb were obtained for Doublon. Detailed information about the optical maps for Anso77 and Doublon is summarized in Table S4. Initial assemblies were hybrid-scaffolded on their respective set of optical maps. No Doublon optical maps were cut during scaffolding, while one Anso77's optical map was cut. However, ten cuts were generated in nine Anso77's contigs, and four cuts were generated in one Doublon's contig. Summary metrics of the hybrid assembly step are summarized in Additional files: Table S4.

We looked for 148 highly conserved SSR markers in the scaffolds (Additional files: Table S2). In Anso77, 21/30 super-scaffolds contained at least two SSR markers while the scaffold 100023, 90 and 100013 contained only one SRR and 6 scaffolds contained none of the 148 markers. The SSR in the 21 scaffolds ordered fully congruently with their order in Harukei genome excepted for two very close SRRs located on chromosome 10. In Doublon, 20/28 contained at least two SSR markers and the scaffold 100023 contained only one. The SSR ordered fully congruently with their position in the Harukei genome excepted for two very close SRRs (same than in Anso77). All scaffolds containing at least two markers were then assigned and spanned. The scaffolds not spanned at this step, one in Doublon and three in Anso77) were less than 4Mb length. They were localized at chromosome extremities, and we assumed to spin them according to Harukei-3 genome using to dot-plot analysis.

Telomeres were identified at both end of two and nine chromosomes, while three and two chromosomes displayed telomeres at one end only in Anso77 and Doublon, respectively.

#### Supplementary Figures

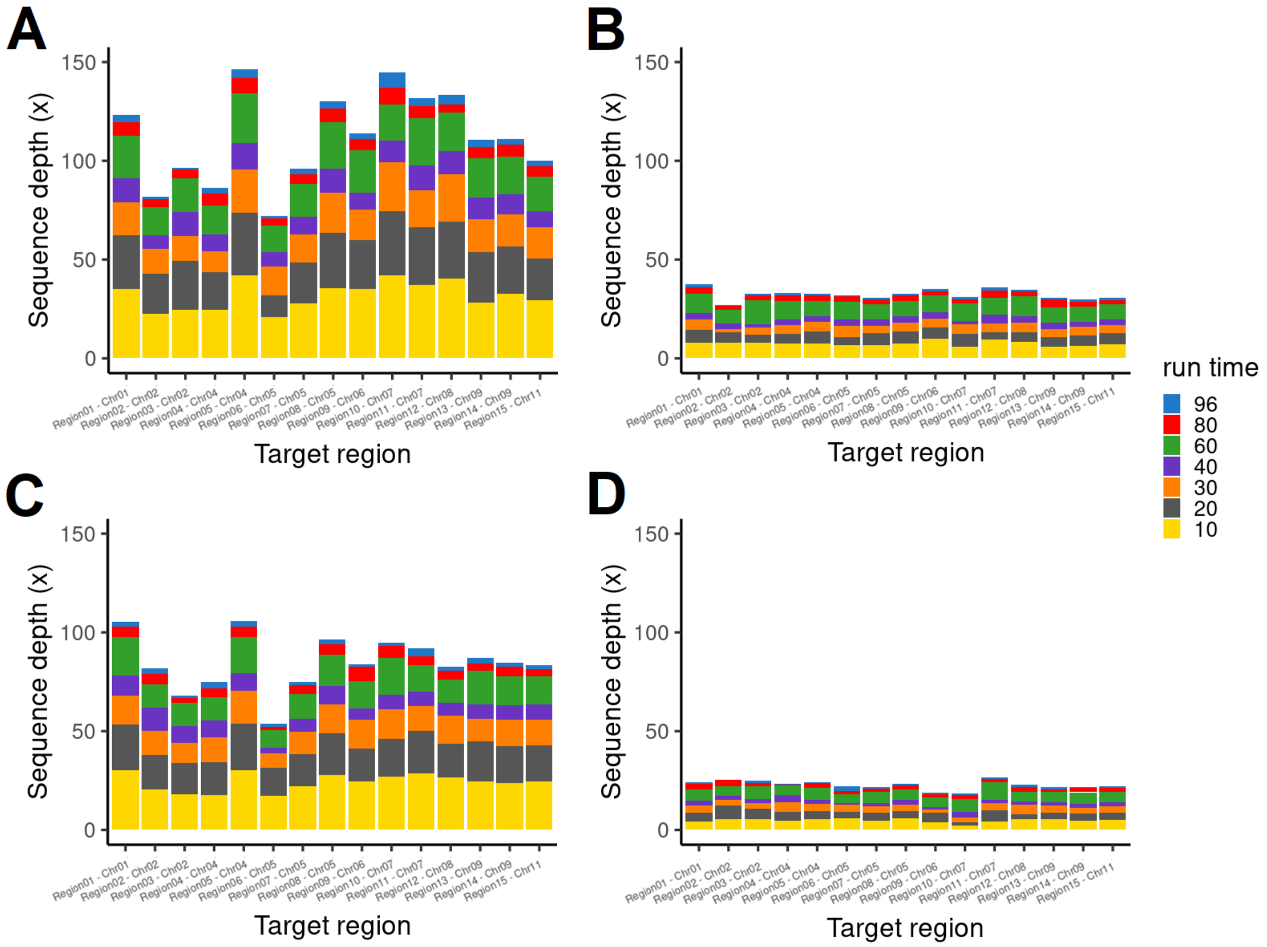


**Figure S1.** Sequence depth on the 15 target regions of Anso77 (A, B) and Doublon (C, D) using NAS (A, C) and a WGS approach (B, D).

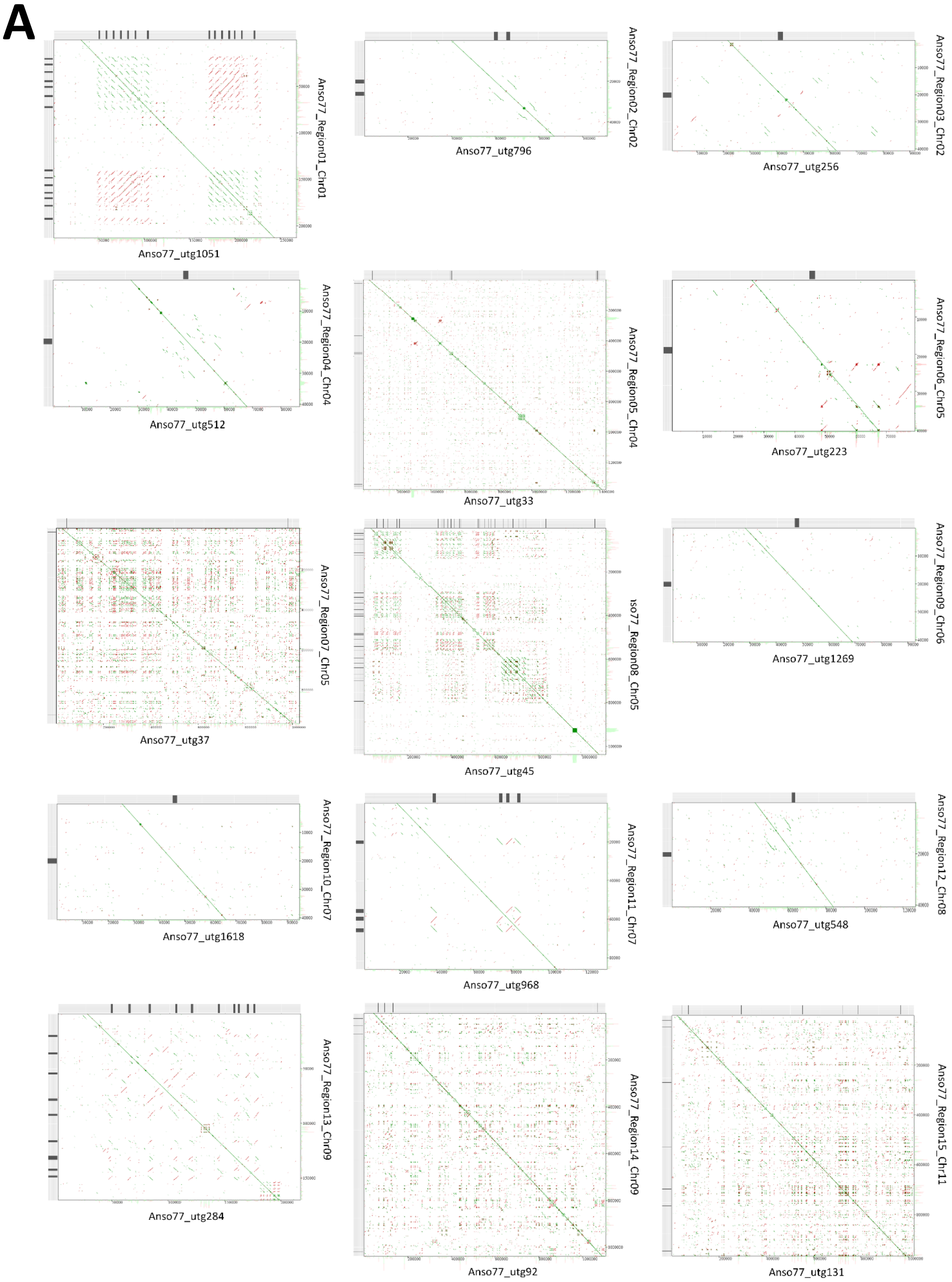
A
Anso77_Region01_Chr01
Anso77_utg1051
Anso77_Region02_Chr02
Anso77_utg796
Anso77_Region03_Chr02
Anso77_utg256
Anso77_Region04_Chr04
Anso77_utg512
Anso77_Region05_Chr04
Anso77_utg33
Anso77_Region06_Chr05
Anso77_utg223
Anso77_Region07_Chr05
Anso77_utg37
so77_Region08_Chr05
Anso77_utg45
Anso77_Region09_Chr06
Anso77_utg1269
Anso77_Region10_Chr07
Anso77_utg1618
Anso77_Region11_Chr07
Anso77_utg968
Anso77_Region12_Chr08
Anso77_utg548
Anso77_Region13_Chr09
Anso77_utg284
Anso77_Region14_Chr09
Anso77_utg92
Anso77_Region15_Chr11
Anso77_utg131

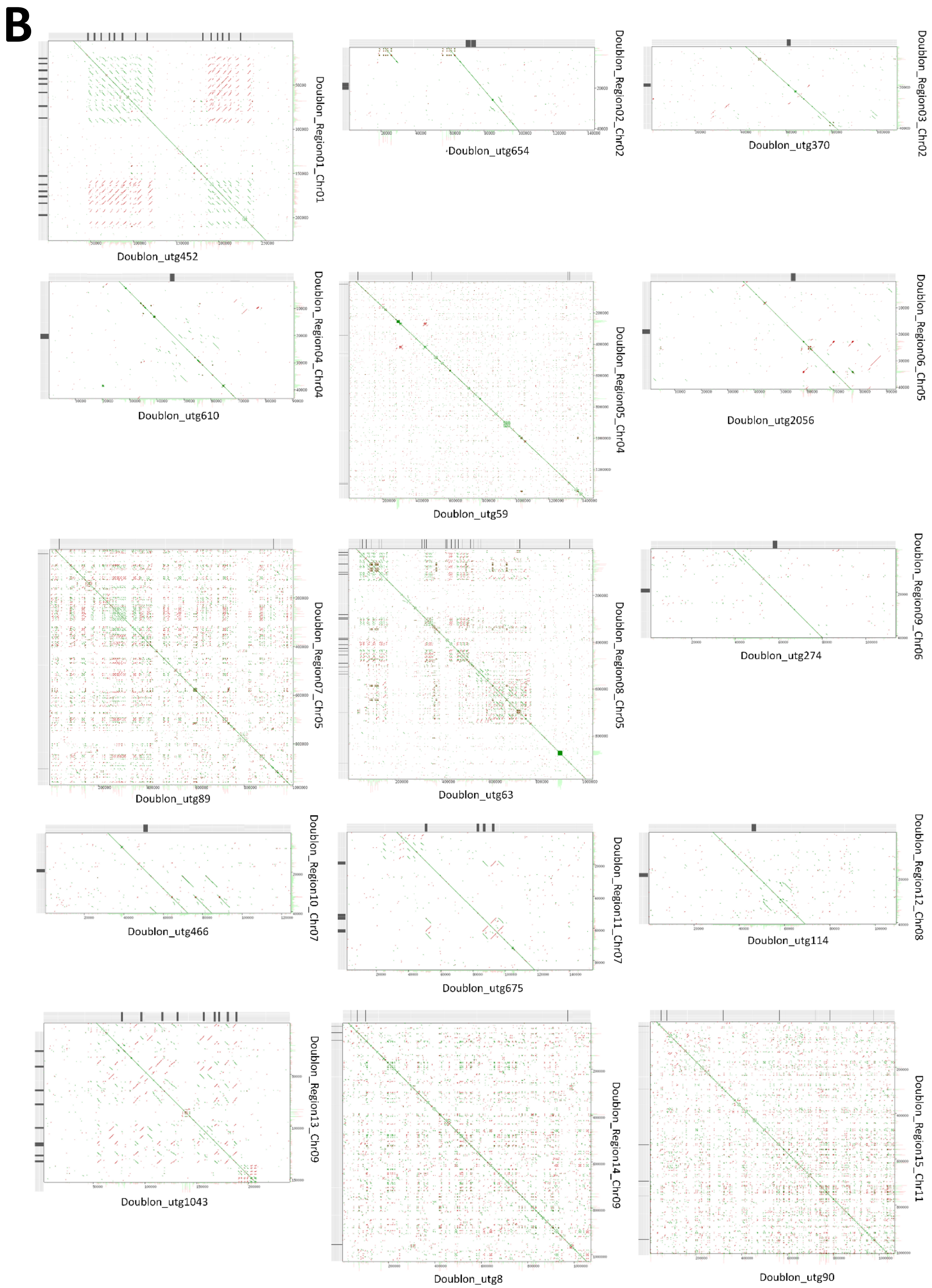


**Figure S2.** Dotplots comparing the NAS assembly (x axis) and reference assembly (y axis) of the 15 target regions of Anso77 (A) and Doublon (B). Positions of the predicted NLR domains are represented with black vertical lines.

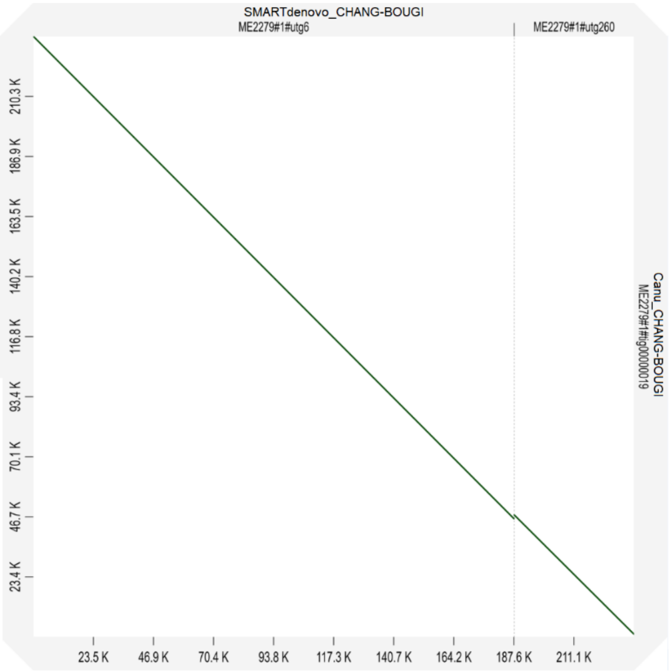


**Figure S3.** Dotplot representing the assembly of the region 13 using SMARTdenovo (x axis) and Canu (y axis). The Canu assembler achieved a contiguous assembly of the region in one single contig.

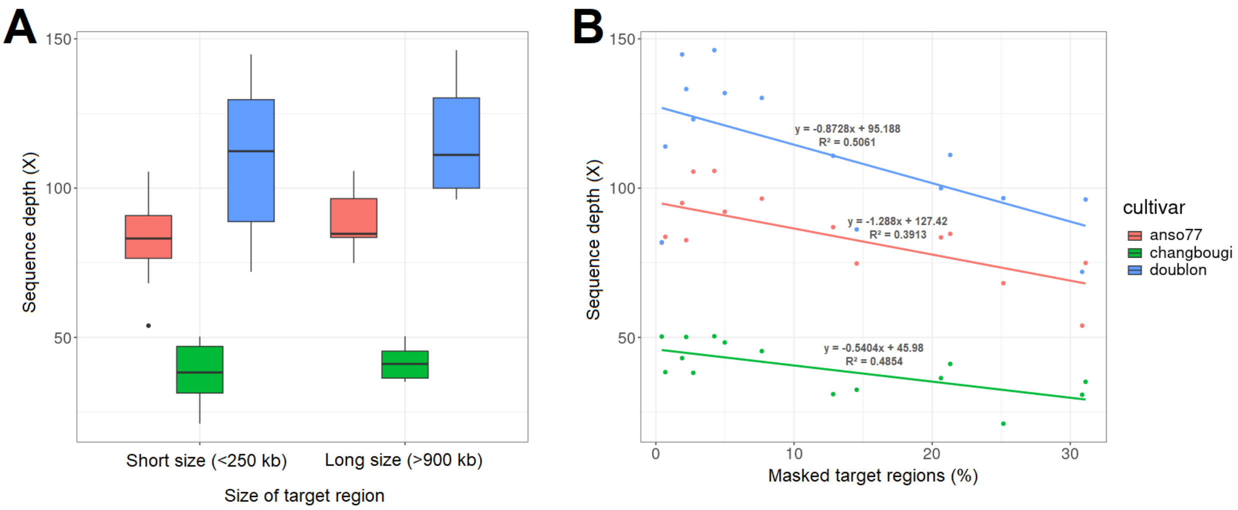


**Figure S4.** Influence of the size of the target regions (A) and the percentage of the target regions masked (B) in the final sequence depth.

### Supplementary Tables

**Table S1 – S2. See Supplementary File 1**

**Table S3.** Summary metrics of the generated reads and Bionano molecules for Anso77 and Doublon.

| | Anso77 | | | | Doublon | | | |
|---|---|---|---|---|---|---|---|---|
| | ONT | 10x | Illumina | Bionano | PacBio | ONT | Illumina | Bionano |
| **Total size (Gb)** | 45.1 | 35.5 | 8.8 | 902 | 40.5 | 2.6 | 7.8 | 805 |
| **Number of reads or molecules* (M)** | 3.8 | 223.0 | 29.0 | 0.4* | 2.8 | 0.21 | 25.69 | 0.3* |
| **Read or molecules* length N50 (kb)** | 29.8 | - | - | 235.8* | 24.7 | 27.3 | - | 256.7* |
| **longest reads (kb)** | 321 | - | - | - | 122 | 253 | - | - |

**Table S4**. Summary metrics of the primary contig assembly, optical mapping and hybrid scaffolding of Anso77 and Doublon.

| | | Anso77 | Doublon |
|---|---|---|---|
| **Contigs** | Number | 159 | 186 |
| | Total size (Mb) | 366.7 | 362.0 |
| | N50 (Mb) | 8.9 | 15.2 |
| | Min. length (Mb) | 0.001 | 0.001 |
| | Max. length (Mb) | 23.6 | 26.9 |
| **Optical maps** | Number | 33 | 28 |
| | Total size (Mb) | 388.8 | 391.3 |
| | N50 (Mb) | 20.5 | 20.6 |
| | Min. length (Mb) | 0.6 | 0.4 |
| | Max. length (Mb) | 30.1 | 30.3 |
| **Hybrid assembly** | Number | 30 | 28 |
| | Total size (Mb) | 365.7 | 359.2 |
| | N50 (Mb) | 19.4 | 19.7 |
| | Min. length (Mb) | 0.4 | 0.2 |
| | Max. length (Mb) | 29.0 | 29.0 |
| | Number of Ns (Mb) | 3.0 | 1.0 |
| **Contigs not scaffolded** | Number | 82 | 113 |
| | Total size (Mb) | 4.0 | 3.9 |
| | N50 (Mb) | 0.1 | 0.06 |
| | Min. length (Mb) | 0.001 | 0.001 |
| | Max. length (Mb) | 0.3 | 0.1 |
